# Purine metabolism regulates DNA repair and therapy resistance in glioblastoma

**DOI:** 10.1101/2020.03.26.010140

**Authors:** Weihua Zhou, Yangyang Yao, Andrew J. Scott, Kari Wilder-Romans, Joseph J. Dresser, Christian K. Werner, Hanshi Sun, Drew Pratt, Peter Sajjakulnukit, Shuang G. Zhao, Mary Davis, Meredith A. Morgan, Alnawaz Rehemtualla, Barbara S. Nelson, Christopher J. Halbrook, Li Zhang, Francesco Gatto, Jianping Xiong, Maria G. Castro, Pedro Lowenstein, Sriram Chandrasekaran, Theodore S. Lawrence, Costas A. Lyssiotis, Daniel R. Wahl

## Abstract

Intratumoral genomic heterogeneity in glioblastoma (GBM) is a barrier to overcoming therapy resistance, and new strategies that are effective independent of genotype are urgently needed. By correlating intracellular metabolite levels with radiation resistance across dozens of genomically-distinct models of GBM, we found that purine metabolites strongly correlated with radiation resistance. Inhibiting purine, but not pyrimidine, synthesis radiosensitized GBM cells and patient-derived neurospheres by impairing DNA repair in a nucleoside-dependent fashion. Likewise, administration of exogenous purine nucleosides protected sensitive GBM models from radiation by promoting DNA repair. Combining an FDA-approved inhibitor of *de novo* purine synthesis with radiation arrested growth in GBM xenograft models and depleted intratumoral guanylates. High expression of the rate-limiting enzyme of *de novo* GTP synthesis was associated with shorter survival in GBM patients. Together, these findings indicate that inhibiting *de novo* purine synthesis may be a promising strategy to overcome therapy resistance in this genomically heterogeneous disease.

## Introduction

Glioblastoma (GBM) is the most common and aggressive adult brain tumor and is associated with profound genomic heterogeneity, which has limited therapy development. Work from The Cancer Genome Atlas (TCGA) and others have defined a diversity of driver alterations in GBM including gene amplifications, mutations, deletions and complex rearrangements of signaling receptors^1,2^. Unfortunately, targeted therapies against these abnormalities have uniformly lacked efficacy in patients with GBM^3,4,5,6^. These disappointing results may be due to the profound *intra-tumoral* genomic heterogeneity that also characterizes GBM. Indeed, single-cell and regional sequencing have shown that the molecular events vary region-to-region and even cell-to-cell within a single GBM^7,8,9^. This heterogeneity may explain why the only therapies that have improved survival in GBMs do not require a precise molecular alteration for activity: radiation (RT), temozolomide, surgery, and tumor treating fields^10^.

RT is a critical treatment modality for GBM patients^11^ and RT-resistance is the primary cause of recurrence and death in GBM. Fewer than 10% of patients with GBM live for 5 years and approximately 80% recur within the high dose radiation field^12, 13^. Thus, efforts to overcome primary RT-resistance are likely to improve outcomes in patients with GBM. Efforts to develop new strategies to overcome RT-resistance have previously used large-scale genomic profiling data to define candidate oncogenic molecular alterations to target in combination with radiation^14, 15, 16^. Due to the profound genomic heterogeneity of GBM, we instead sought to define therapeutic strategies that could overcome RT-resistance independently of genotype.

Altered metabolism is a hallmark of cancers including GBM, is regulated by cell-intrinsic and -extrinsic factors, and could potentially regulate therapy resistance independently of genotype^17, 18, 19, 20, 21^. Importantly, disparate oncogenic alterations can activate common metabolic pathways, such as glycolysis^22^. Thus, GBMs with profound intra-tumoral genomic heterogeneity may have relatively common metabolic phenotypes that in turn mediate resistance to radiation. We therefore sought to determine how metabolism mediates RT-resistance in a genotype-independent fashion to identify new therapeutic targets and biomarkers for GBM.

## Results

### Nucleotide metabolites correlate with RT-resistance in GBM

To determine the characteristics of RT-resistance in GBM, we performed clonogenic survival assays on 23 distinct immortalized GBM cell lines and found a wide distribution of intrinsic RT sensitivities (Fig. 1A). These lines were chosen because they had been genomically profiled by the Cancer Cell Line Encyclopedia (CCLE), were publicly available from cell line repositories (ATCC, DSMZ and JCRB) and were amenable to both reproducible metabolomic analysis and the clonogenic survival assay in a uniform media (DMEM). None had mutations in isocitrate dehydrogenase (IDH) 1 and thus were models of primary GBM. Intrinsic RT-resistance did not correlate with cell proliferation rate (Fig. S1A) or cell cycle distribution (Fig. S1B). Ionizing radiation causes double-stranded breaks (DSBs) in DNA and rapidly results in the phosphorylation of histone H2A variant H2AX, which can be readily detected by immunoblot, flow cytometry or immunofluorescence^23^. Both RT-resistant (U87 MG and A172) and -sensitive cell lines (KS-1 and U118 MG) had high levels of γ-H2AX staining at 30 min and 2 h after RT (Fig. 1B). However, γ-H2AX staining returned to baseline by 24 h after RT in the RT-resistant lines, while it remained persistently elevated in the RT-sensitive lines. Thus, RT-resistance in this GBM cell line panel is associated with an ability to effectively repair RT-induced DSBs.

**Figure 1.**
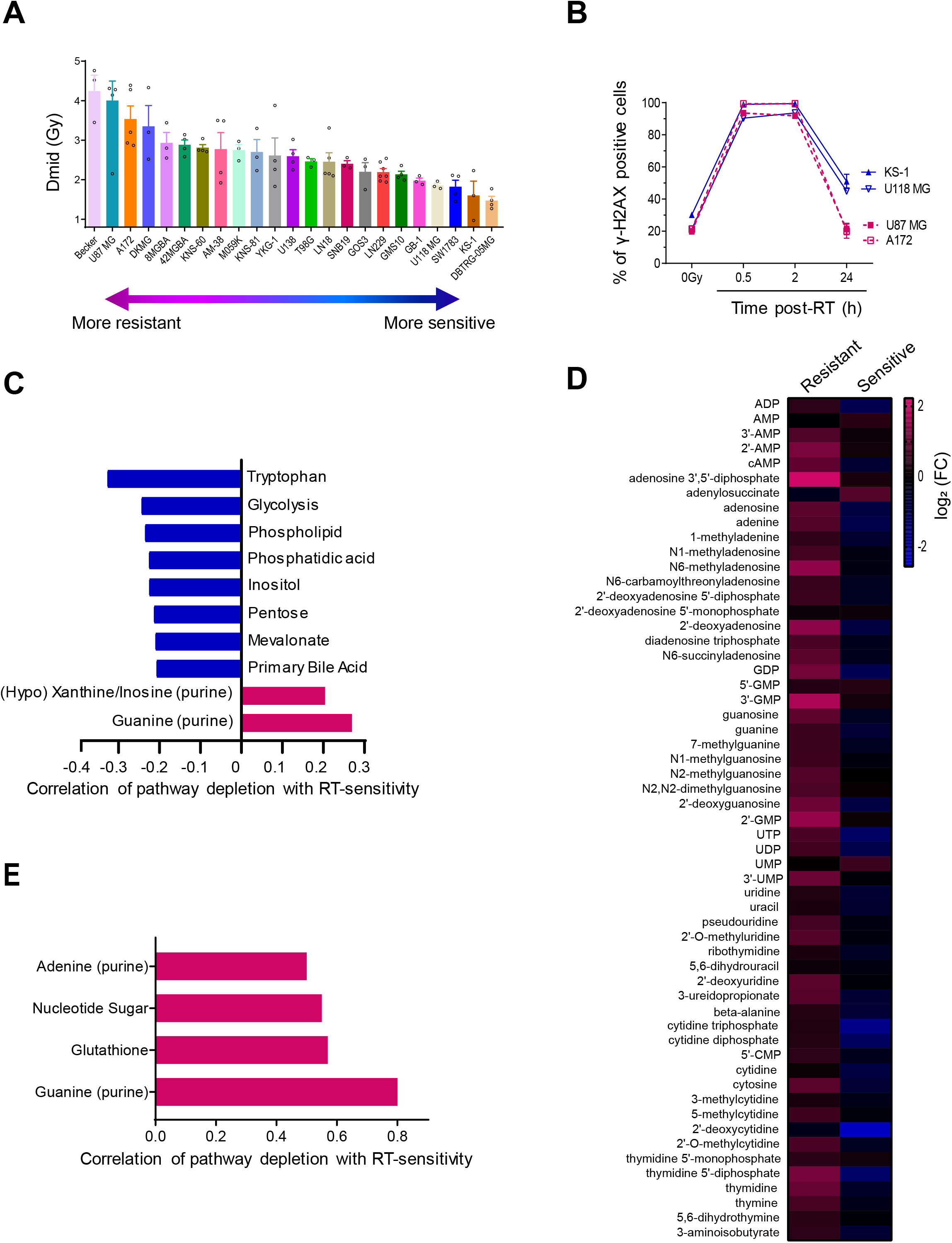
Increased levels of nucleobase-containing metabolites are associated with RT-resistance in GBM. **(A)** Indicated GBM cell lines were plated at clonal density, and colonies were counted 10 to 14 days later. **(B)** RT-resistant (U87 MG and A172) and RT-sensitive (KS-1 and U118 MG) GBM cell lines were irradiated with 8 Gy, followed by γ-H2AX flow cytometry analysis at 0 h, 0.5 h, 2 h, or 24 h following RT. **(C)** Targeted metabolomics analysis on 23 GBM cell lines. Metabolites were grouped in to corresponding pathways and an average pathway-level correlation with RT-sensitivity was determined. Pathways with downregulated metabolites that were significantly correlated with RT-sensitivity are shown. **(D)** RT-resistant and -sensitive GBM cell lines were irradiated with 8 Gy, and harvested 2 h after RT and analyzed by targeted LC-MS/MS. Red panels indicate the increase of metabolites and green panels indicate the depletion of metabolites comparing to the metabolite levels in the parental GBM cells. **(E)** Pathways with downregulated metabolites post-RT that are significantly correlated with RT-sensitivity are shown (Pearson’s correlation; *p* < 0.05).

Our group and others have postulated that abnormal metabolism in GBM may cause RT-resistance^24, 25^. Using molecular data obtained from the CCLE, we asked whether transcript expression of metabolic enzymes could predict for GBM RT-resistance. Consistent with our prior work^24^, increased expression of *IDH1* was associated with GBM RT-resistance (Fig. S1C), presumably because this enzyme is an important source of NADPH in GBM. Glutamine synthetase (*GLUL*) expression was also associated with RT-resistance (Fig. S1D), consistent with the prior reports^26^. *IDH3a*, a critical mediator of oxidative ATP production through the TCA cycle, was instead associated with RT-sensitivity (Fig. S1E). Gene set enrichment analysis revealed that three out of the top 10 most associated gene sets with RT-sensitivity were related to oxidative ATP production (Fig. S1F). No such metabolic gene sets were found among the top 10 associated gene sets with RT-resistance (Fig S1G). This relative lack of actionable metabolic targets suggested a need to measure metabolism itself rather than the levels of metabolism-related transcripts.

We therefore performed targeted metabolomic analysis on each of the 23 GBM cell lines during unperturbed exponential growth (These data accompany this manuscript as a supplementary file). Metabolites were grouped into corresponding pathways and correlations between pathway-level changes and RT-resistance were determined to identify metabolic phenotypes associated with RT resistance. All determinations of both RT-resistance and unperturbed metabolome were performed in consistent cell culture media. Downregulation of metabolites involved in *de novo* purine synthesis (inosinates and guanylates) were positively correlated with RT-sensitivity (*p* < 0.03) (Fig. 1C&S1H). Downregulation of the cytidine pathway was the third most-correlated metabolic pathway with RT-sensitivity, but was not statistically significant (*p* = 0.08). Thus, GBMs with lower nucleotide pools, especially purines, were more likely to be RT-sensitive.

We then asked how purine metabolism changed after cells were exposed to RT. Two hours after RT, a time point when DNA damage had occurred (Fig. 1B) but cells had not yet arrested or died (Fig. S1B), both purine and pyrimidine metabolites increased in RT-resistant cell lines (Fig. 1D). RT-sensitive cell lines, however, increased neither purines nor pyrimidines following RT (Fig. 1D). In the post-RT setting, depleted guanylates were again the metabolic feature most correlated with RT-sensitivity (*p* = 0.0001) (Fig. 1E). This analysis also revealed that decreased levels of metabolites related to glutathione, the primary cellular antioxidant, were significantly associated with RT-sensitivity (*p* = 0.02), which is consistent with the well-known oxidative mechanism by which radiation kills cells^27^ Depletion of adenylates, the other main purine species, were also significantly associated with RT-sensitivity (Fig. 1E). Together, these results suggested that high levels of nucleobase-containing metabolites, especially purines, were related to GBM RT-resistance.

### Supplementing nucleotide pools protects GBMs from radiation by facilitating DSB repair

We next sought to determine whether the relationship between high nucleobase-containing metabolites and RT-resistance in GBM was causal. Nucleotide-poor RT-sensitive GBM cell lines were supplemented with cell-permeable nucleosides (adenosine, guanosine, cytidine, thymidine and uridine) and RT-sensitivity was determined by clonogenic assay (Fig. 2A). RT-sensitive GBM cell lines (U118 MG, DBTRG-05MG, and GB-1; Fig. 1A) were protected from RT by exogenous nucleosides with enhancement ratios (ER) ranging between 0.6 and 0.8 (Figs. 2B-D). The RT-protection conferred by nucleosides was associated with decreased RT-induced DSBs. Indeed, in all three sensitive cell lines, RT alone caused a peak of γ-H2AX foci within 30 min that did not return to baseline by 24 h (Figs. 2E-G). Treatment with exogenous nucleosides decreased γ-H2AX foci at 0.5, 2, 6 and 24 h following RT (Figs. 2E-G; Figs. S2A-C) and reduced the DSBs presented 24 h after RT to near baseline levels.

**Figure 2.**
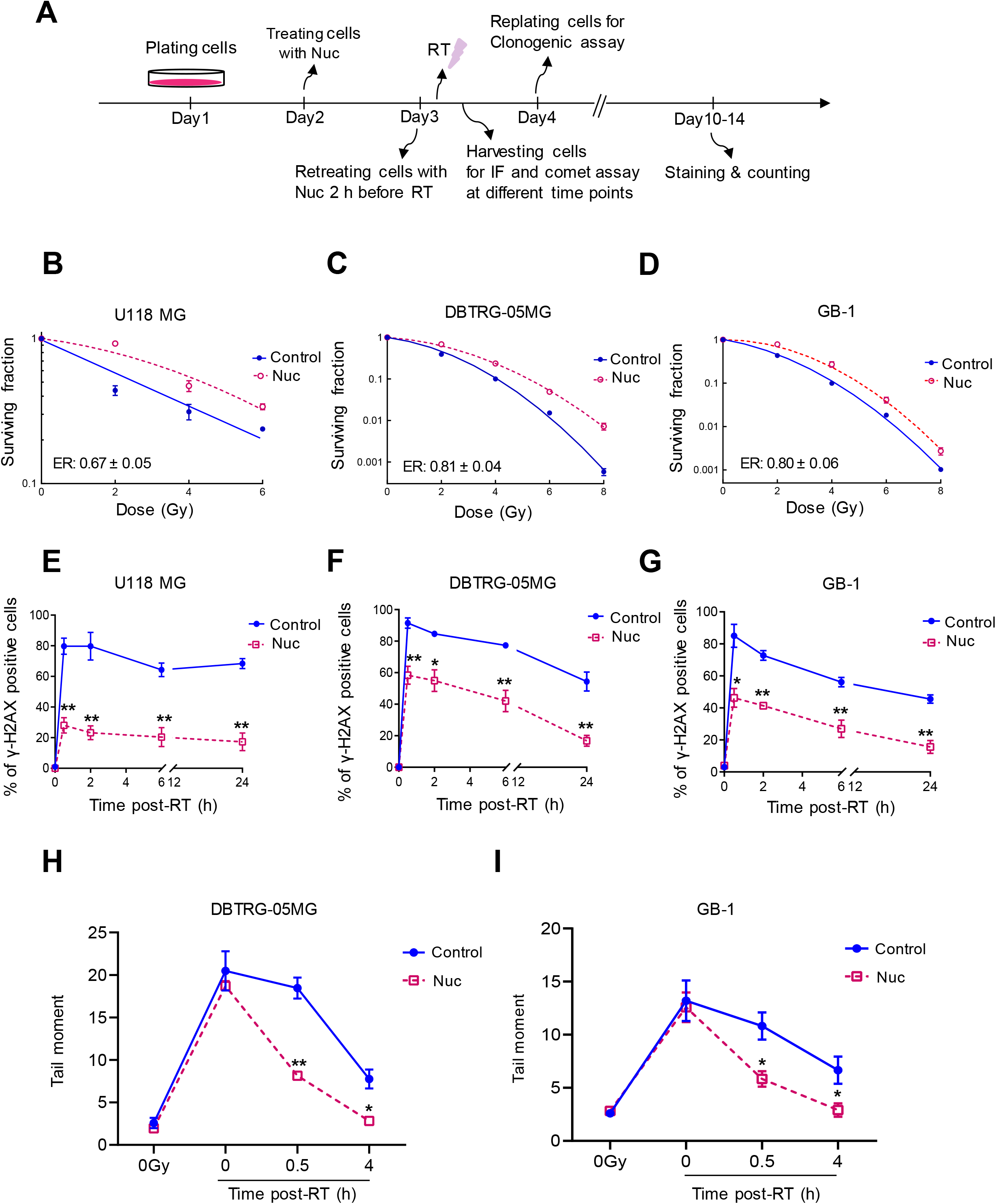
Supplementing nucleotide pools promotes DNA repair and RT-resistance in GBM. **(A)** A schematic timeline of treatment in the RT-sensitive cell lines. U118 MG, DBTRG-05MG, or GB-1 cells were treated with exogenous nucleoside pools (8 x) for 24 h, and retreated with nucleosides 2 h before RT with indicated doses, followed by IF, comet assay, or clonogenic assay. **(B-D)** U118 MG, DBTRG-05MG, and GB-1 cells were treated as described in Fig. 2A, and plated at clonal density 24 h post-RT, and colonies were counted 10 to 14 days later. ER indicates Enhancement Ratio. **(E-G)** Cells were treated as discussed above, and harvested for γ-H2AX foci staining at indicated time point post-RT. **(H&I)** DBTRG-05MG or GB-1 cells were treated as discussed above, and cells were harvested at different time points for alkaline comet assay. Note: cells were irradiated and harvested on ice for the 0 h time point (4 Gy; 0h). *, *p* < 0.05 and **, *p* < 0.01, compared with control; error bars indicate SEM from 3 to 5 biologic replicates.

Because exogenous nucleosides reduced γ-H2AX staining both early and late after RT, this assay did not allow us to determine whether nucleosides were preventing the induction of RT-induced DSBs or facilitating their repair. We therefore performed the alkaline comet assay^28^, which measures physical DNA damage and can distinguish between changes in DSB induction and early DSB repair. GBM cells were irradiated on ice and harvested immediately post-RT (4 Gy; 0 h), which eliminates DNA repair, or incubated at 37 °C and harvested at differing times post-RT (4 Gy; 0.5 h, 4 h). Nucleosides did not change the amount of DNA damage induced when cells were irradiated on ice and harvested immediately (Fig. 2H&I). However, exogenous nucleosides decreased the RT-induced DNA damage that was present after repair was allowed to proceed for 0.5 and 4 h in two RT-sensitive GBM cell lines, DBTRG-05MG (*p* < 0.01 for 0.5 h; *p* < 0.05 for 4 h) and GB-1 (*p* < 0.05 for 0.5 h and 4 h; Figs. 2H&I; Figs. S2D&E). These results indicate that supplementing nucleotide pools in RT-sensitive GBMs is sufficient to facilitate the repair of RT-induced DSB.

### Inhibition of *de novo* purine synthesis slows DSB repair and radiosensitizes RT-resistant immortalized GBM cell lines

Based on the above data, we next asked if lowering nucleotide pools would slow DSB repair and radiosensitize RT-resistant models of GBM. We chose to inhibit *de novo* GTP synthesis (Fig. 3A) because guanylates were the metabolic pathway most correlated with RT-resistance and most GBMs are thought to rely on *de novo* nucleotide synthesis rather than nucleotide salvage^29^. Drugs inhibiting *de novo* GTP synthesis, such as mycophenolic acid (MPA) and its orally bioavailable prodrug mycophenolate mofetil (MMF), are FDA-approved to treat immune-mediated disorders and are being investigated as anticancer therapeutics^30^. Treatment with a clinically-relevant concentration of MPA (10 μM)^31^ reduced GTP levels by more than 10 fold, increased inosine monophosphate levels by more than 10 fold and slightly increased ATP levels, consistent with inhibition of inosine monophosphate dehydrogenase (IMPDH), the molecular target of MPA and the rate limiting step in *de novo* GTP synthesis (Figs. 3A-D). We investigated the effects of MPA on two RT-resistant cell lines (Fig S3A) and found that MPA radiosensitized both in a concentration-dependent fashion (Figs 3E&F). ER ranged from 1.2 in U87 MG (column 1 vs 2; *p* < 0.01) and 1.3 in A172 (column 1 vs 2;*p* < 0.01) at 1 μM MPA treatment and 1.7 in U87 MG (column 1 vs 3; *p* < 0.0001) and 1.98 (column 1 vs 3; *p* < 0.001) in A172 at 10 μM MPA, which is comparable to the radiosensitizing effects of temozolomide, the standard radiosensitizer used in GBM^32^. The radiosensitizing effects of MPA were abrogated when cells were co-treated with exogenous nucleosides (column 3 vs 5; ER: 1.7 vs 1.0; *p* < 0.01 in U87 MG and 2.3 vs 1.2; *p* < 0.05 in A172; Figs. 3E&F), indicating that MPA exerted its radiosensitizing effects through nucleotide depletion rather than off-target effects. Unlike in RT-sensitive GBM cell lines, exogenous nucleosides did not further protect RT-resistant GBM cell lines from RT (column 1 vs 4; Figs. 3E&F). We speculate that RT-resistant cells are already rich in nucleotides (Figs. 1C&D), which limits the ability of further nucleoside supplementation to further protect cells.

**Figure 3.**
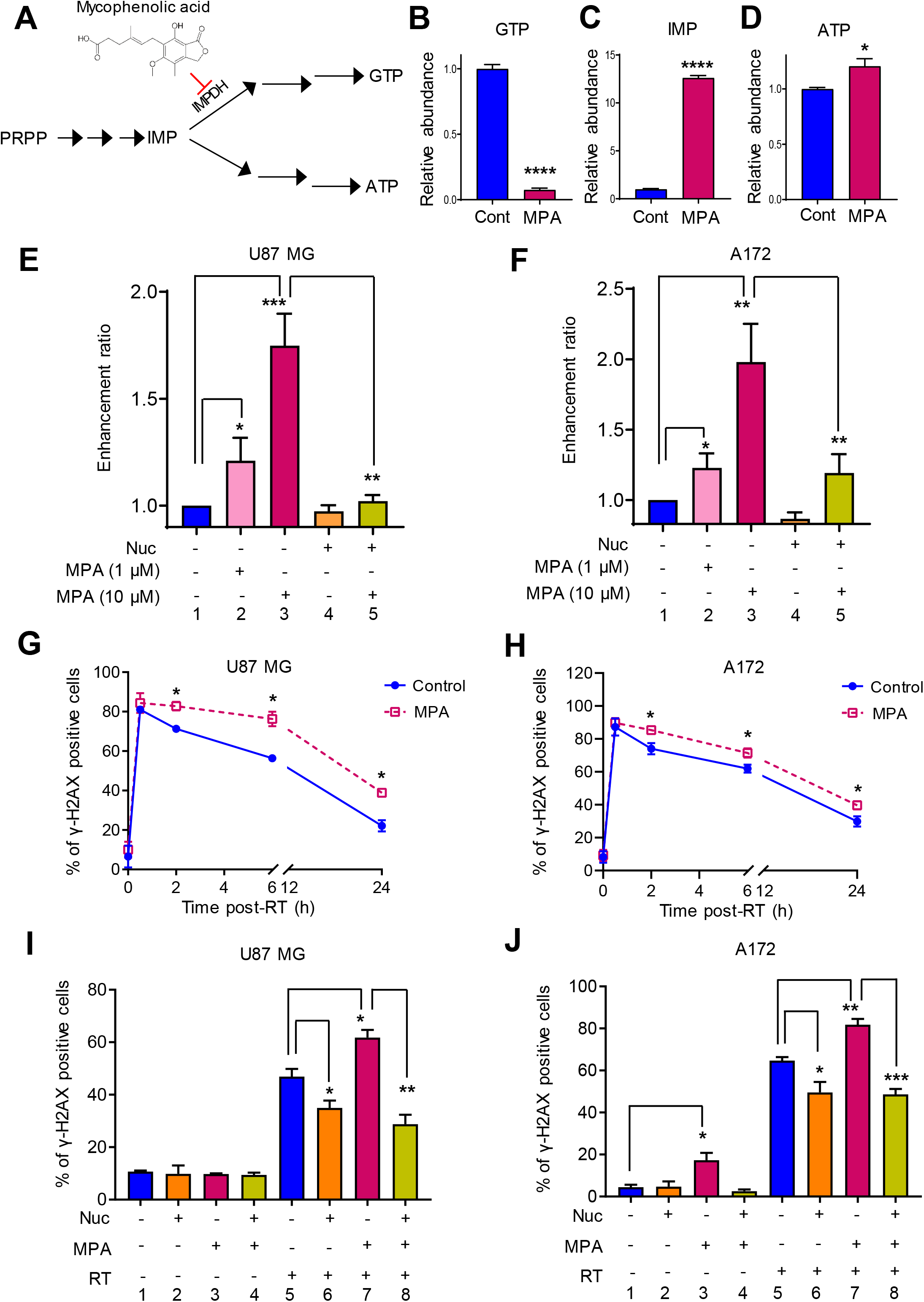
Inhibiting *de novo* purine synthesis impairs DNA repair and radiosensitizes RT-resistant GBM cells. **(A)** A schematic diagram of *de novo* purine biosynthesis. PRPP: Phosphoribosyl pyrophosphate; IMP: inosine monophosphate; GTP: Guanosine-5’-triphosphate; ATP: Adenosine triphosphate; IMPDH: Inosine-5’-monophosphate dehydrogenase; MPA: mycophenolic acid; **(B-D)** U87 MG cells were treated with MPA at 10 μM for 24 h, and then harvested and analyzed by targeted LC-MS/MS. *,*p* < 0.05, and ****,*p* < 0.0001 compared to control. **(E&F)** After treatment with indicated conditions, cells were replated for colonogenic assay and colonies were stained and counted 10 to 14 days later. **(G&H)** Cells were treated with different doses of MPA for 24 h and then irradiated with 4 Gy, and cells were harvested at indicated time point for γ-H2AX foci staining. **(I&J)** After treatment with indicated conditions, cells were harvested and fixed for IF γ-H2AX foci staining 6 h post-RT. Note: Figs. D-I, *, *p* < 0.05; **, *p* < 0.01, and ***, *p* < 0.001, compared with control; error bars indicate SEM from 3 to 5 biologic replicates.

We reasoned that GTP depletion may sensitize GBMs to RT by slowing DSB repair, much as nucleoside supplementation promoted DSB repair. Consistent with this hypothesis, the combination of MPA and RT increased γ-H2AX foci at various time points compared to RT alone in both U87 MG and A172 cells (*p* < 0.01; Figs. 3G&H; Figs. S3B&C). This increase was rescued by the administration of exogenous nucleosides (column 7 vs 8; MPA vs MPA + Nuc, *p* < 0.01 in U87 MG and *p* < 0.001 in A172; Figs. 3I&J; Figs. S3D&E). Thus, inhibition of *de novo* purine synthesis radiosensitizes RT-resistant GBM cell lines by slowing DSBs repair in a nucleoside-dependent fashion.

### Inhibition of *de novo* purine synthesis radiosensitizes primary patient-derived GBM neurospheres

The above findings were obtained in GBM cell lines that we found to be resistant to RT in our initial profiling. While tractable for metabolomic and clonogenic survival assays, such immortalized GBM models may not fully recapitulate the histopathologic or molecular features of GBM tumors in patients^33^. To overcome these limitations, we used primary patient-derived GBM neurosphere lines, referred to as HF2303 and MSP12^34^, to confirm our findings. These primary GBM cells form neurospheres when grown in serum-free conditions, are inherently resistant to irradiation and are thought to represent the cellular subtypes that mediate GBM recurrence after therapy^35, 36^.

Because neurospheres are not amenable to the clonogenic survival assay, we instead performed a long-term viability assay to assess the effects of radiation. Primary neurospheres were treated with MPA, and/or nucleosides, followed by various doses of RT and then replated as single cells and allowed to grow for 7-10 days before viability was assessed. Compared to the control, treatment with MPA increased the sensitivity of GBM neurospheres to RT (ER: 1.4 ± 0.1 for HF2303 and 1.7 ± 0.2 for MSP12). MPA-induced radiosensitization could be reversed when it was combined with exogenous nucleosides (ER: 1.03 ± 0.04 for HF2303 and 0.9 ± 0.2 for MSP12) (Figs. 4A&B).

**Figure 4.**
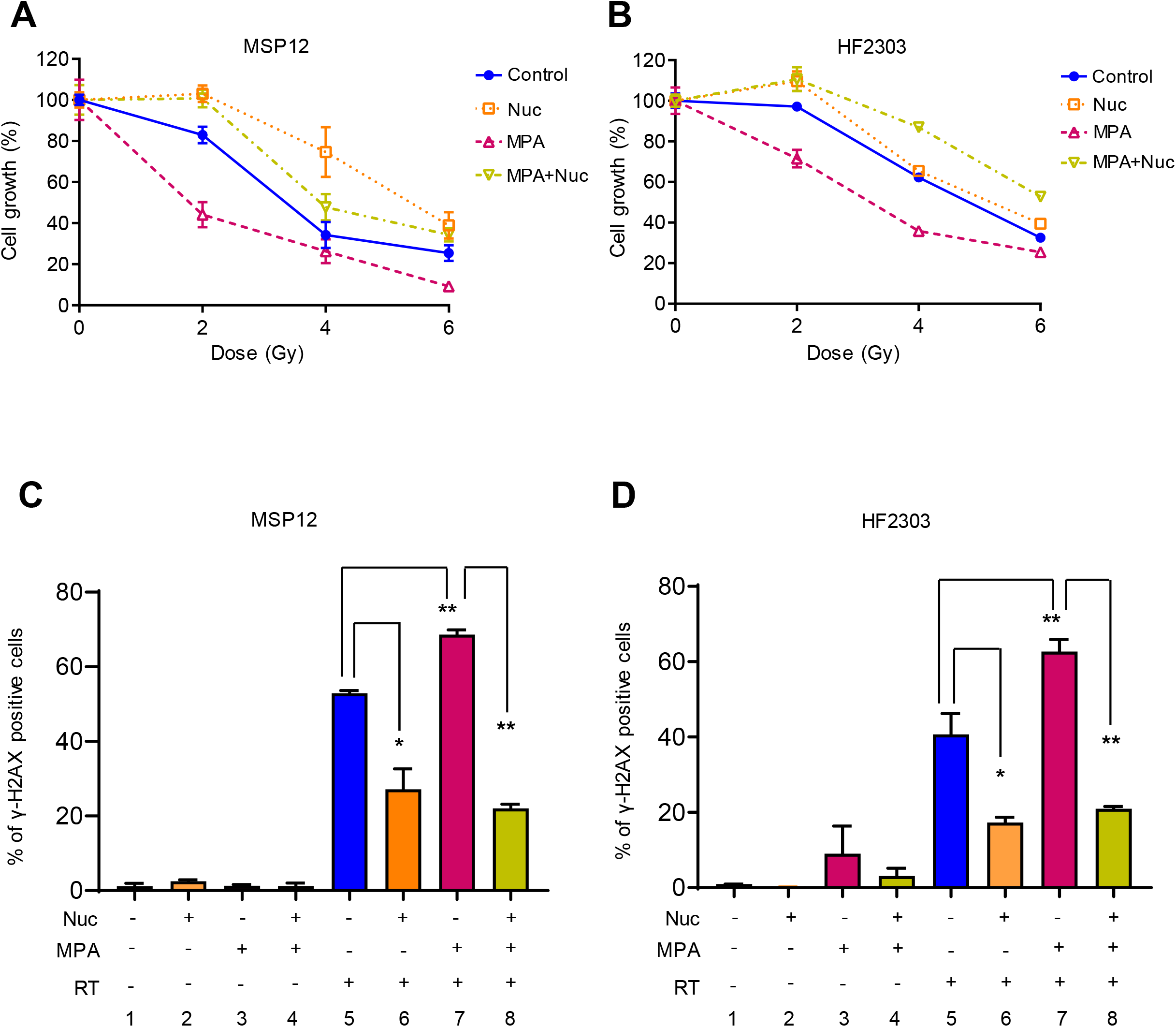
Inhibiting *de novo* purine synthesis radiosensitizes primary patient-derived GBM neurospheres in a nucleoside-dependent fashion. **(A-D)** MSP12 or HF2303 neurospheres were treated as discussed above, and replated to the 96-well plate (2000 cells/well) for the CelltiterGlo assay 24 h post-RT **(A&B)**, or fixed for IF γ-H2AX foci staining 6 h post-RT **(C&D)**. *, *p* < 0.05 and **, *p* < 0.01, compared with control; error bars indicate SEM from 3 to 5 biologic replicates. Note: Fig. A and B are representative figures from 3 repeated experiments.

Furthermore, γ-H2AX foci formation assay confirmed that MPA increased RT-sensitivity in primary neurosphere models by enhancing DSBs (column 5 vs 7; *p* < 0.01), which were reversed by exogenous nucleoside treatment (column 7 vs 8; *p* < 0.01; Figs. 4C&D; Figs. S4A&B). Hence, inhibition of *de novo* purine synthesis radiosensitizes GBM in a nucleoside-dependent fashion in primary patient-derived models of GBM.

### Purines, not pyrimidines, are the dominant nucleotide species that govern RT-resistance and DNA repair in GBM

We next sought to understand more precisely which nucleotide species were mediating RT-resistance and DNA repair in GBM. We took advantage of teriflunomide, which is an FDA-approved inhibitor of dihydroorotate dehydrogenase (DHODH), the rate limiting enzyme in *de novo* pyrimidine synthesis (Fig. S5A). Teriflunomide is used to treat multiple sclerosis^37^ and is under investigation as an anticancer drug^38^. Here, we found that inhibition of *de novo* pyrimidine synthesis by teriflunomide decreased the levels of pyrimidines in GBM cells (Figs. S5B-F). However, teriflunomide treatment did not overcome RT-resistance in RT-resistant GBM cell lines (Figs. 5A&B), nor did it impair the ability of these cell lines to repair RT-induced DSBs as measured by γ-H2AX foci (Figs. 5C&D; Figs. S5G&H).

**Figure 5.**
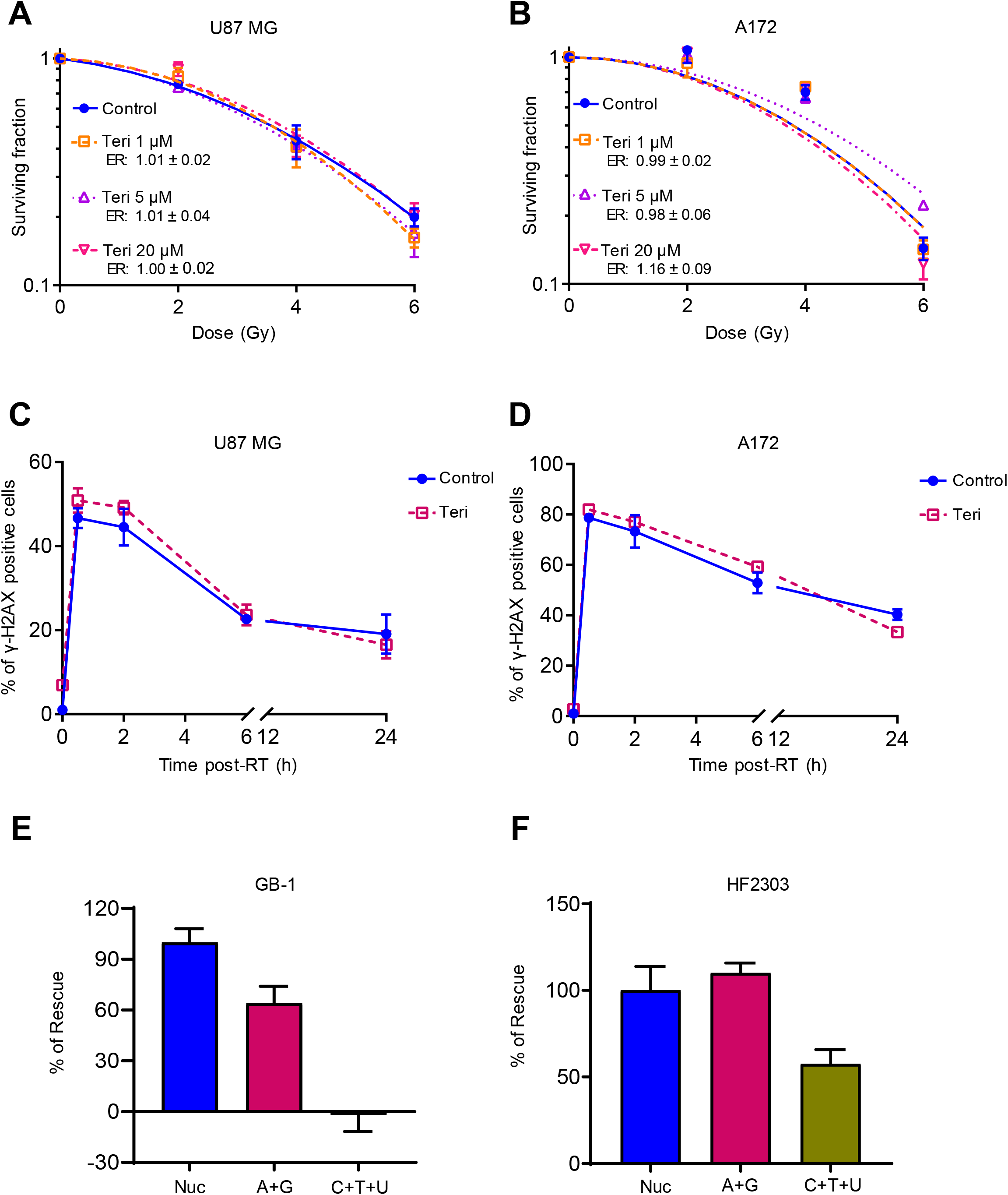
Modulating pyrimidine pools has minimal effects on DNA repair and RT-resistance in GBM. **(A&B)** U87 MG and A172 cells were treated with varying doses of teriflunomide for 24 h, and then irradiated. Cells were replated for colonogenic assay 24 h post-RT. **(C&D)** Cells were treated with different doses of teriflunomide for 24 h and then irradiated with 4 Gy, and cells were harvested at indicated time point post-RT for γ-H2AX foci staining. Error bars, SEM from 3 biologic replicates. **(E&F)** Cells were treated with exogenous nucleoside pools (8 x), or a combination of Adenosine + Guanosine (A + G), or Uridine + Cytidine + Thymidine (U + C + T) for 24 h, and retreated with indicated nucleosides 2 h before RT (4 Gy), followed by IF γ-H2AX foci staining 6 h post-RT.

Pyrimidine supplementation also failed to fully protect GBM cell lines and neurospheres from radiation. In the RT-sensitive GB-1 cell line, purines alone (adenosine and guanosine) promoted the repair of RT-induced DSBs nearly as much as pooled nucleosides. Pyrimidines alone (cytidine, uridine and thymidine), did not promote the repair of RT-induced DSBs (Figs. 5E&S5I). In the HF2303 patient-derived neurosphere model, purines stimulated DNA repair as much as pooled nucleosides (Figs. 5F&S5J). Pyrimidines alone failed to fully recapitulate the effects of pooled nucleosides. Thus, purines appear to play a greater role in mediating DNA repair and RT-resistance in GBM than do pyrimidines.

### MMF slows GBM tumor and is augmented by combination with RT

To further confirm these *in vitro* findings, we established flank xenografts using a U87 MG immortalized cell line (Fig. 6) and determined whether combining RT and inhibition of *de novo* GTP synthesis had similar beneficial anti-GBM effects *in vivo*. We utilized mycophenolate mofetil (MMF), the orally bioavailable pro-drug of MPA that is FDA-approved to treat organ rejection. Once xenografts were 80 to 100 mm^3^ in size, the mice were randomized into two groups: one for endpoint studies and a second to monitor pharmacodynamics response (Fig. 6A). We found that numerous guanylates increased immediately following RT, and this increase was abrogated when RT was combined with MMF (Fig. 6B). The protein level of γ-H2AX increased post-RT, and increased further when combined with MMF (Fig. 6C). MMF by itself caused little γ-H2AX staining.

**Figure 6.**
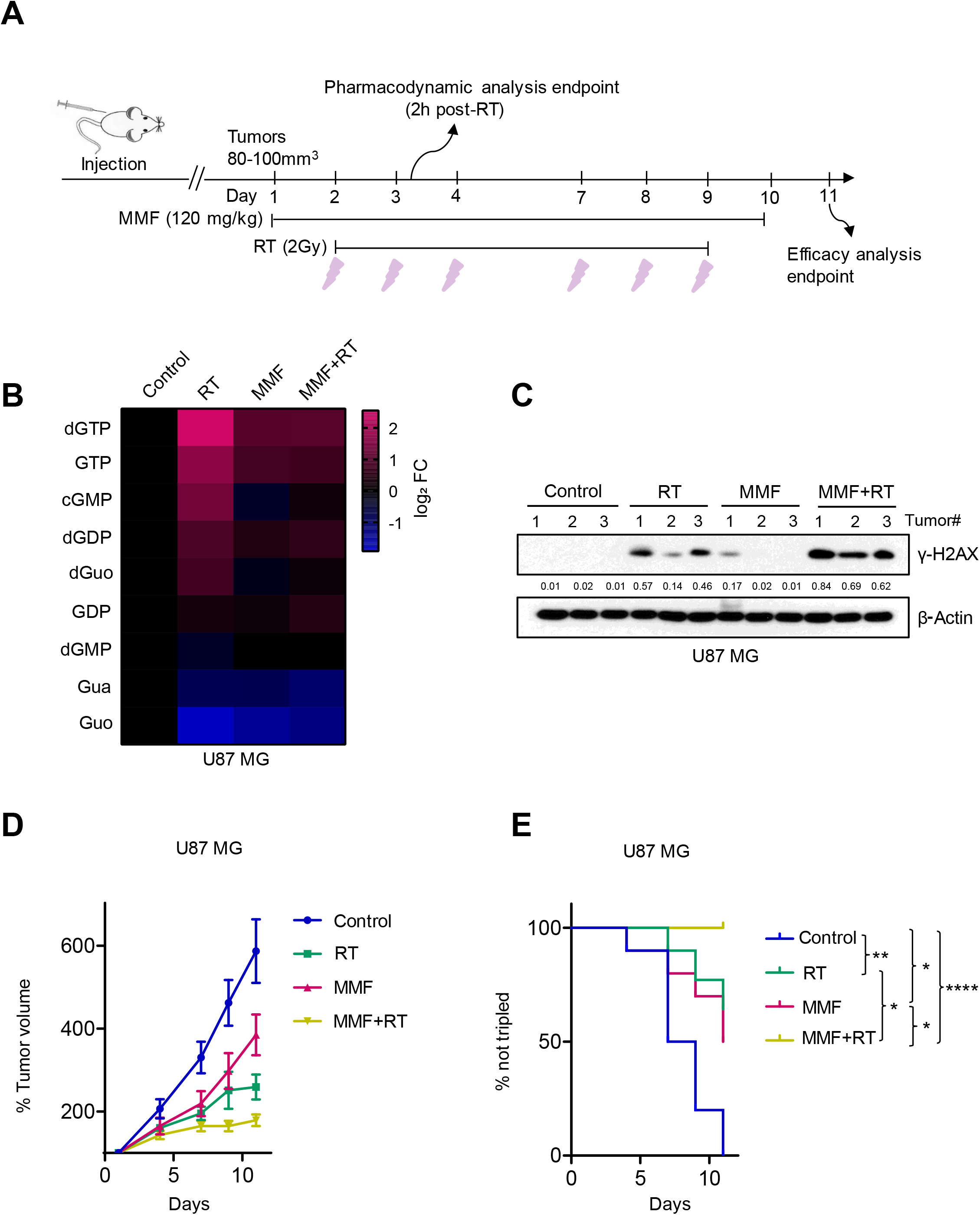
MMF augments RT efficacy against immortalized GBM xenografts. **(A)** A schematic timeline of U87 MG flank model. U87 MG cells (2 x 10^6^) were injected into flanks and tumors were allowed to form. Mice were randomly divided and treated as described in Material and Methods. **(B)** Flash-frozen tumors harvested 2 h post-RT in biological group were analyzed by LC-MS/MS, sample sizes for the 4 groups are 3 (Control), 6 (RT), 3 (MMF) and 5 (RT + MMF). **(C)** Tumors from each group were ground and lysed for immunoblotting assay with indicated antibodies. The bands were quantified using Image J software and the quantified numbers were labeled under each band. **(D)** Tumor volumes for the indicated treatment subgroups of the efficacy group are normalized to the individual tumor sizes defined on day 1. Error bars indicate SEM from 10 tumors from 5 mice per group. **(E)** Kaplan-Meier estimates of time to tumor tripling. *, *p* < 0.05; **, *p* < 0.01; ****, *p* < 0.0001.

Treatment with RT or MMF alone had a modest but significant effect on tumor growth compared with untreated tumors. The combination of RT and MMF nearly arrested tumor growth (Fig. 6D). Furthermore, combined RT and MMF treatment significantly increased the time to tumor tripling compared with single RT (*p* < 0.05), single MMF (*p* < 0.05), or no treatment (*p* < 0.0001; Fig. 6E). IHC staining confirmed that combined RT and MMF treatment showed the lowest expression of proliferation marker Ki-67 compared to the other three groups (Figs. S6A&B).

### MMF slows primary patient-derived GBM xenograft tumor growth *in vivo*

To further confirm the data obtained in immortalized GBM xenografts, we established xenograft flank models using two primary patient-derived neurospheres (HF2303 and MSP12). The mice were randomized as in the immortalized xenograft model (Figs. 7A&S7A). Consistent with the immortalized xenograft model, several guanylates were elevated 2 h following RT in HF2303 xenografts. This increase was again abrogated when MMF was administered along with RT (Fig. 7B). γ-H2AX protein level increased after RT and further increased when MMF was combined with RT in both the HF2303 and MSP12 models (Figs. 7C&S7C). In both HF2303 and MSP12 tumors, single agent MMF and RT modestly slowed tumor growth. However, combined MMF and RT significantly slowed tumor growth (Figs. 7D&S7D), and increased the time to tumor tripling (Figs. 7E&S7E). Median days to tumor tripling are 12 (Control), 16 (MMF), 23 (RT), and 33 (RT + MMF) for HF2303 and 10 (Control), 12 (MMF), 13.5 (RT), and 19 (RT + MMF) for MSP12, respectively. Consistent with this efficacy, combined MMF and RT decreased the expression of the cell proliferation marker Ki-67 combined to either treatment in isolation in both the MSP12 and the HF2303 models (Fig. S7G). The effects of treatments on normal tissues was minimal, as reflected by relatively unchanged body weight during drug treatment (Figs. S7B&F).

**Figure 7.**
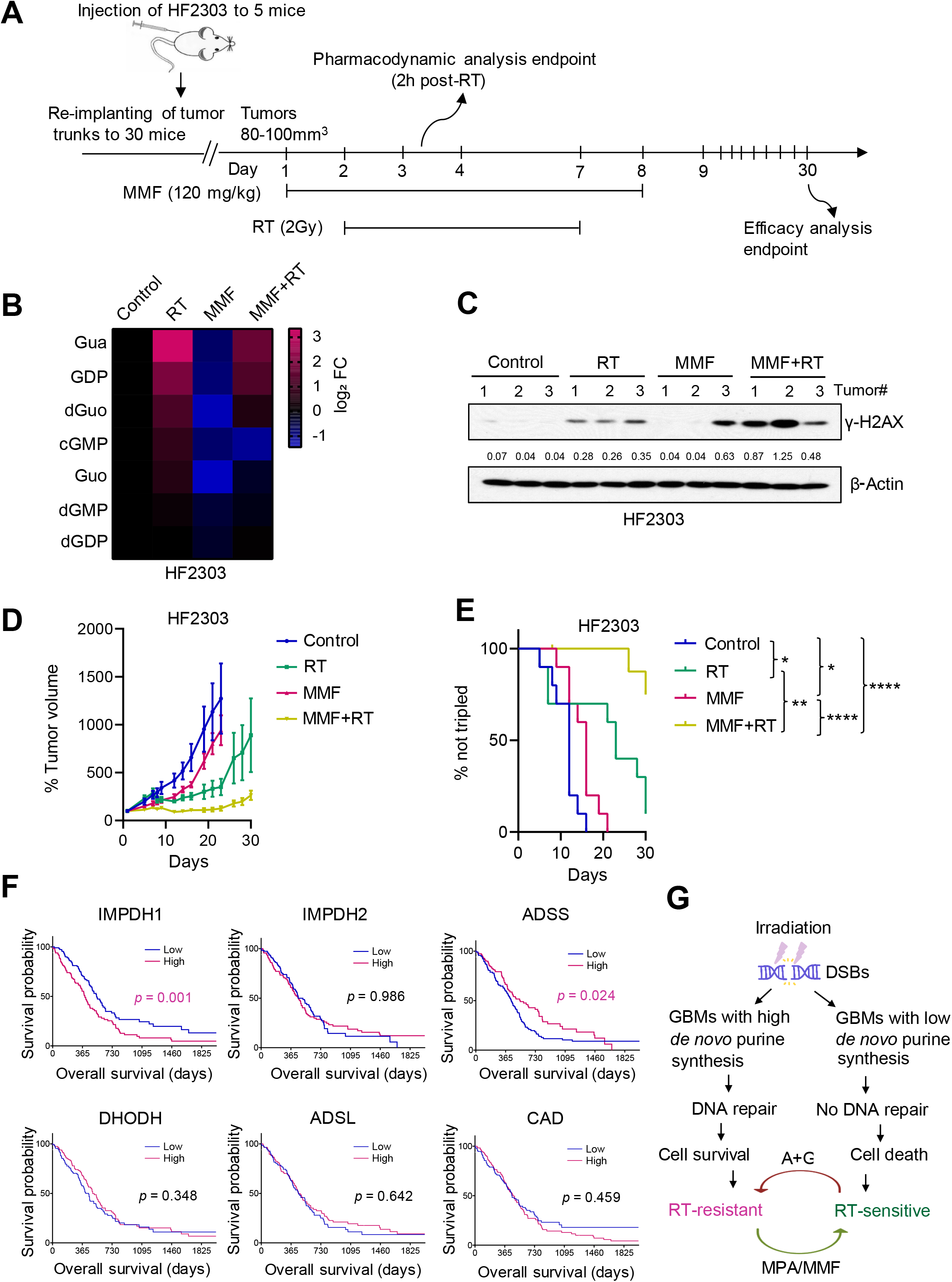
MMF augments RT efficacy against patient-derived GBM xenografts and associations between patient survival and expression of genes involved in nucleotide synthesis. **(A)** A schematic timeline of HF2303 flank model. HF2303 xenograft GBM model was established and treated as described in the Material and Methods. **(B)** Flash-frozen tumors harvested 2 h post-RT in biological group were analyzed by MS. **(C)** Tumors from each group were ground and lysed for immunoblotting assay with indicated antibodies. The bands were quantified using Image J software and the quantified numbers were labeled under each band. **(D)** Tumor volumes for the indicated arms of the efficacy group are normalized to the individual tumor sizes defined on day 1. Error bars indicate SEM from 10 tumors from 5 mice per group. **(E)** Kaplan-Meier estimates of time to tumor tripling. *, *p* < 0.05; **, *p* < 0.01; ****, *p* < 0.0001. (**F**) The mRNA expression of the rate-limiting enzymes of nucleotide pathways in 235 patients from the Pan-Cancer Atlas with newly diagnosed IDHwtGBM and the survival analysis. (**G**) Working model. RT induces DSBs in GBMs. High *de novo* purine synthesis promotes GBM survival by stimulating dsDNA repair, cell survival and recurrence after RT. Supplementing cells with purines (A+G) promotes RT-resistance while inhibiting *de novo* purine synthesis with mycophenolic acid (MPA) or mycophenolate mofetil (MMF) promotes RT-sensitivity.

### High expression of *IMPDH1* is associated with inferior survival in patients with molecularly-defined aggressive brain tumors

Finally, we asked if these data were reflected in the outcomes of patients with brain tumors. We identified 235 patients from the Cancer Genome Atlas Pan-Cancer Atlas with newly diagnosed primary *IDH1* wild type gliomas (so-called “molecular GBMs”) whose samples had passed the Pan-Cancer quality assurance, the vast majority of whom received RT^39^. Increased transcript expression (> median) of *IMPDH1*, the rate limiting enzyme in *de novo* GTP synthesis and target of MPA/MMF, was associated with inferior overall survival (HR 0.59, 95% CI 0.43-0.81, *p* = 0.001). Increased expression of *ADSS* and *ADSL*, the rate limiting enzymes in *de novo* ATP synthesis, were not associated with worse survival. Furthermore, increased expression of rate-limiting enzymes in *de novo* pyrimidine synthesis (*DHODH* and *CAD*) was not associated with decreased survival in patients with newly diagnosed IDHwt glioma (Fig. 7F). Together, these *in vitro, in vivo* and patient-level data suggest that purines, especially GTP, mediate RT-resistance and DNA repair in GBM and that inhibition of *de novo* GTP synthesis could be a promising therapeutic strategy for GBM, especially when combined with RT.

## Discussion

Intratumoral genomic heterogeneity in GBM has limited the efficacy of personalized targeted therapies. To overcome this barrier, we sought to discover metabolic pathways that caused RT-resistance in GBM independently of genotype. By analyzing how intracellular metabolite levels correlated with GBM RT resistance, we found that low levels of nucleobase-containing metabolites were strongly associated with sensitivity to RT. This association was causal, as supplementing GBM cells with exogenous nucleosides protected them from RT by promoting the repair of RT-induced DSBs in DNA. The protective effects of these nucleosides were primarily due to purines rather than pyrimidines. We also showed that this relationship between nucleotide pools and RT-resistance could have therapeutic applications. Depleting intracellular GTP pools with an FDA-approved inhibitor of *de novo* purine synthesis sensitized GBM cell lines to RT by slowing the repair of dsDNA breaks. Depleting pyrimidine pools had no such effects. These results were recapitulated in patient-derived neurosphere models of GBM and in several *in vivo* xenograft models when MMF was given in combination with radiation. Additionally, high expression of the rate limiting enzyme in *de novo* GTP synthesis was associated with inferior survival in patients with molecularlydefined aggressive gliomas regardless of grade, while high expression of the rate limiting enzymes in *de novo* ATP synthesis or pyrimidine synthesis was not. In summary, we have found that *de novo* purine synthesis is a targetable vulnerability that causes RT-resistance in GBM and is associated with inferior patient survival (Fig. 7G).

These results add to a growing body of literature indicating that *de novo* purine synthesis contributes to the aggressive behavior of GBM and other cancers^40^. High rates of *de novo* purine and pyrimidine synthesis promote the maintenance and tumorigenic capacity of glioma-initiating cells, which are thought to contribute to therapy resistance and tumor recurrence in GBM^41, 42^. Our data suggests that the high rates of *de novo* purine synthesis in these tumorigenic cells may be directly related to their enhanced ability to repair RT-induced DNA damage and mediate tumor recurrence. *De novo* purine synthesis can generate both GTP and ATP. *De novo* GTP synthesis is preferentially upregulated in GBM, while *de novo* ATP synthesis is similarly active in both normal brain tissue and GBM. This upregulation of GTP synthesis promotes tRNA and rRNA synthesis, nucleolar transformation and GBM proliferation^43^. This importance of GTP was recapitulated in our studies, as guanylates were most strongly correlated with GBM RT-resistance and inhibiting *de novo* GTP synthesis alone was sufficient to overcome GBM RT resistance. These findings suggest that the FDA-approved inhibitors of *de novo* GTP synthesis currently used to treat inflammatory diseases should be evaluated for therapeutic benefit in patients with GBM, particularly in combination with RT. Because normal glia and neural stem cells primarily relay on guanylate salvage rather than *de novo* synthesis^43^, such a therapeutic strategy may have minimal normal tissue toxicity.

Like the small number of therapies with proven benefit in GBM, inhibitors of *de novo* GTP synthesis do not require a precise oncogenic event for activity. Therefore, these inhibitors may have clinical benefit despite the intratumoral genomic heterogeneity that characterizes GBM. Indeed, many of the heterogeneous oncogenic alterations that drive GBM including mutations, deletions or amplifications in *PTEN, EGFR* and *PIK3CA* can cause the similar metabolic phenotype of elevated *de novo* purine synthesis^18, 19, 20, 44^. Thus, a genomically heterogenous GBM^7,8,9^ may exhibit a relatively homogeneous metabolic phenotype of elevated *de novo* purine synthesis, which could be exploited therapeutically to overcome RT-resistance. Here, we observed this in our heterogeneous PDX flank models (Fig. 7 & S7).

Our study raises several questions. MMF has efficacy when combined with RT in numerous flank models of GBM. Whether MMF is similarly efficacious for GBMs growing in their natural intracranial environment is unknown. Fortunately, this drug has a low molecular weight, is relatively lipophilic and accumulates to efficacious concentrations in normal mouse brain^45^. All of these factors suggest that MMF will have favorable intracranial activity, but its ability to accumulate in GBM and deplete intratumoral GTP levels in human patients with GBM should be empirically tested. How purines, especially GTP, regulate RT-resistance and dsDNA repair in GBM remains to be defined. Because modulating pyrimidine levels did not cause similar effects as modulating purines, we believe that this link is likely due to active signaling, perhaps through a GTP-activated protein, rather than the more simplistic explanation that modulating nucleotide pools alters the availability of the substrates for DNA repair. Our experiments were entirely carried out in models of GBM without mutations in *IDH1* or *IDH2*, which represent the vast majority of GBMs in patients. Whether the rarer secondary GBMs containing the *IDH1* or *IDH2* mutation exhibit a similar relationship between *de novo* purine synthesis and therapy resistance remains to be defined.

In summary, we have defined *de novo* purine synthesis as the dominant metabolic pathway that mediates RT-resistance in GBM. These findings have motivated the development of a clinical trial to test whether MMF achieves effective concentrations in GBM tissue in patients and whether it is safe and effective in combination with RT for patients with this disease.

## Methods

### Cell culture and reagents

The details of the origins of the GBM cell lines used in our study are listed in supplemental Table 1. HF2303 primary neurosphere, which was originally described by Dr. Tom Mikkelsen at Henry Ford Hospital (Detroit, MI)^36^, was a kind gift from Dr. Alnawaz Rehemtulla and MSP12 was a gift from Drs. Pedro Lowenstein and Maria Castro. Cell line authentication was performed by the originating cell line repositories and then used immediately upon receipt. Cell lines were re-authenticated using short tandem repeat profiling if they had been using for longer than 1 year. Mycoplasma test (Cat# LT07-418, Lonza) were performed monthly in our lab. All the immortalized GBM adherent cells were cultured in DMEM (Cat# 11965-092, Gibco) supplemented with 10% FBS (Cat# S11550, ATLANTA biologicals), 100 μg/mL Normocin (Cat# ant-nr-1, InvivoGen) and 100 U/mL Penicillin-Streptomycin-Glutamine (Cat# 10378-016, Gibco). Primary patient-derived GBM neurospheres (HF2303 and MSP12) were cultured in DMEM-F12 (Cat# 10565-018, Gibco) supplemented with B-27 supplement (Cat# 17504-044, Thermofisher), N2 supplement (Cat# 17502-048, Thermofisher), 100 U/mL Penicillin-Streptomycin (Cat# 15140122, Thermofisher), 100 μg/mL Normocin (Cat# ant-nr-1, InvivoGen), and 20 ng/mL epidermal and fibroblast growth factors (Cat# AF-100-15 and #100-18B, PeproTech). Individual nucleosides, including uridine (U3003), thymidine (T1895), cytidine (C4654), adenosine (A4036), guanosine (G6264), teriflunomide (SML0939), and MPA (M5255), were purchased from Sigma. The other reagents used in the study include Acummax (Cat# AM-105, Innovative cell technologies Inc), Teriflunomide (Cat# SML0936, Sigma), MPA (Cat # M5255, Sigma), MMF (Cat# S1501, Selleckchem), and pooled nucleosides (Cat# ES-008-D, Millipore).

### Immunofluorescence

Immortalized or primary GBM cells were plated and treated with indicated conditions. Cells were then collected and fixed at indicated time points using 4% paraformaldehyde. Primary neurospheres were mixed with the Histogel (Cat# HG-4000-012, Thermo Scientific) and transferred to tissue cassettes for paraffin embedding, section and staining. The γ-H2AX foci were detected with mouse monoclonal antibody anti-phospho-Histone H2AX (Cat #05-636, Millipore) and goat anti-mouse IgG cross-adsorbed secondary antibody, alexa fluor 594 (Cat# A-11005, Invitrogen). DNA was stained with DAPI. γ-H2AX foci were scored for each experimental arm in more than 100 cells of immortalized GBM cells and more than 15 spheres of primary GBM cells. The foci threshold, which is used to define a positive cell, was 10 for immortalized cells and 3 for primary GBM neurospheres.

### Flow cytometry analysis

For cell cycle analysis, cells were harvested and fixed with pre-cooled 70% ethanol overnight, followed by staining with propidium iodide/RNase A staining buffer (Cat# 550825, BD Bioscience) in the dark at room temperature for 15 min as described previously^46^. γ-H2AX analysis was performed as previously described^47^. Briefly, samples were incubated with a mouse monoclonal antibody anti-phospho-Histone γ-H2AX (Cat #05-636, Millipore) and with a FITC conjugated anti-mouse secondary antibody, followed by staining with propidium iodide/RNase A to assess total DNA content. Data were analyzed on a FACScan flow cytometer (Becton Dickinson) with FlowJo software (Tree Star).

### Clonogenic survival assay

Clonogenic assays were performed as previously described^24, 48^. Cells were irradiated with indicated doses and treated with or without compounds, following by replating at clonal density. Plates were stained and colonies containing > 50 cells were counted after 10 to 14 days of growth. The RT enhancement ratios were calculated as the ratio of the mean inactivation dose (Dmid) under control conditions divided by the mean inactivation dose under drug-treated conditions. Dmid is defined as the mean inactivating dose of radiation^49^. For sphere-forming assays, enhancement ratios were calculated as the ratio of the GI50 in the control condition divided by the GI50 in the treated condition.

### Alkaline Comet assay

GB-1 and DBTRG-05MG cells were plated and treated with indicated conditions at different time points. Single-cell gel electrophoretic comet assays were performed under alkaline conditions according to the previous report^28^. Briefly, cells were combined with 1% LM Agarose (Cat# IB70051, IBI SCIENTIFIC) at 40 °C at a ratio of 1:10 (vol/vol) and immediately pipetted onto slides. For cellular lysis, the slides were immersed in lysis solution (Cat# 4250-050-01, R&D SYSTEMS) overnight at 4 °C in the dark, followed by washing in alkaline unwinding solution (200 mM NaOH, 1 mM EDTA, pH > 13) for 30 min. Then, the slides were immersed in alkaline electrophoresis solution (200 mM NaOH, 1 mM EDTA, pH > 13) to be subjected to electrophoresis at 21 V for 30 min and stained in 2.5 μg/mL propidium iodide (Cat# P3566, Invitrogen) for 20 min. All images were taken with a fluorescence microscope and analyzed by Comet Assay IV software (Perceptive Instruments). For quantification, the tail moment, a measure of both amount and distribution of DNA in the tail, was used as an indicator of DNA damage. Comet-positive cells were scored in random fields of cells, with more than 100 cells for each experimental condition.

### Celltiter Glo cell viability assay

MSP12 or HF2303 primary GBM cells were dissociated and plated into 6-well plates and allowed to form spheres. 3-4 days post-plating, neurospheres were treated with nucleosides and/or MPA, followed by irradiation. Twenty-four hours after RT, cells were disassociated into single cells and replated into 96-well plates with around 2000 cells per well. After growing for 7 to 10 days, cell viabilities were detected using CellTiter-Glo^®^ 3D Reagent (Cat# G9682, Promega) following the manufacturer’s protocol.

### *In vivo* xenograft models

All mouse experiments were approved by the University Committee on Use and Care of Animals at the University of Michigan. C.B-17 SCID mice (female, 4-7 weeks old) were obtained from Envigo and maintained in specific pathogen-free conditions. For U87 MG xenograft model (Fig. 6A), 2 x 10^6^ cells were resuspended in 1:1 PBS:Matrigel (BD Biosciences) and subcutaneously injected into the bilateral dorsal flanks of 30 mice. For HF2303 (Fig. 7A) and MSP12 (Fig. S7A) primary neurosphere xenograft models, 6 x 10^6^ (HF2303) or 2 x 10^6^ (MSP12) were first injected into the bilateral dorsal flanks of 5 mice for each model and when the tumor volume was around 500-600 mm^3^, tumors were harvested and cut into small pieces of similar size and implanted into bilateral flanks of 30 mice for each model. For the three xenograft models, once the tumor volume reached ~100 mm^3^ after injection or implanting, mice were randomized into two major groups (biological and efficacy groups), which were further subdivided into four arms, including vehicle control (0.5(w/v) methylcellulose/0.1% (v/v) Polysorbate 80), MMF alone, RT alone, or combined RT and MMF. MMF (120 mg/kg) was dissolved in vehicle and administered via oral gavage once daily 2 h before radiation. Radiation (2 Gy/fraction) was administered over 6 (U87 MG) or 4 (HF2303 and MSP12) daily fractions on weekdays using a Philips RT250 (Kimtron Medical) as previously described^24^ The tumor volume and body weight were measured three times weekly. Tumor volumes were determined using digital calipers and the formula (π/6) (Length × Width^2^).

### Immunoblotting assay

Xenografts were ground and lysed using RIPA lysis buffer (Cat# 89900, Thermo Scientific) supplemented with PhosSTOP phosphatase inhibitor (Cat# 04906845001, Roche) and complete protease inhibitor tablets (Cat# 1187358001Roche) as described previously^50^. Proteins were detected with antibodies of γ-H2AX (Cat# 05-636, Millipore) and β-actin (Cat# 4967, Cell Signaling Technology).

### Immunohistochemical staining

Mouse tumors were harvested, fixed in 10% formalin, and embedded in paraffin. Immunohistochemical analyses were performed using the ABC Vectastain Kit (Vector Laboratories) as previously described^51^. After deparaffinization, rehydration, antigen retrieval, and blocking, the tumor tissue slides were incubated with primary Ki-67 antibody (Cat# 550609, BD Biosciences) at 4°C overnight.

### CCLE and Gene Set Enrichment Analysis

Relationships between radiation resistance and transcript levels were determined across 23 cell lines using simple least-squares linear regression between empirically determined Dmid (Fig. 1A) and gene expression as determined by the Broad Institute Cancer Cell Line Encyclopedia (CCLE) version 2012-10-18^52^. For genes of interest (*IDH1, IDH3a* and *GLUL*), a t-test on the slope coefficient was run to check for statistical significance. P-values were adjusted for multiple testing using the Benjamini-Hochberg correction. Genes were then rank-ordered by correlation coefficient and gene set enrichment analysis (GSEA, Figs. S1F-G) was performed to identify the top gene sets correlated with RT-resistance and RT-sensitivity^53^.

### Mass spectrometry sample preparation

GBM cells subjected to indicated conditions were washed once with 1x PBS and quenched at indicated time points with ice-cold 80% methanol on dry ice. Parallel plates were used for protein concentration and subsequent normalization. The methanol-cell mixtures were scraped off plates and transferred to 1.5 mL conical tubes. After centrifuging at maximum speed for 10 min, samples were normalized by protein concentration and stored at −80°C. Samples were dried by speedvac and resuspended in 50:50 Methanol/H_2_O for LC-MS/MS analysis. GBM tumor samples were homogenized by physical disruption in cold (−80°C) 80% methanol. Fractions were clarified by centrifugation, normalized to tissue weights and then lyophilized by speed vac. Dried pellets were resuspended in 1:1 methanol: H2O before LC-MS/MS analysis.

Samples were run in triplicate on an Agilent QQQ 6470 LC-MS/MS with ion pairing chromatography acquiring dynamic multiple reaction monitoring (dMRM) for 226 metabolites with a delta retention time (RT) window of one minute. Data was preprocessed by applying a threshold area of 3,000 ion counts and a coefficient of variation (CV) of 0.5 among triplicates. Metabolites falling below and above these thresholds, respectively, were then manually inspected for peak integration. All chromatography analysis was done with Agilent MassHunter Quantitative Analysis 9.0.647.0.

Initial cell metabolic profiling shown in Figs. 1C-E were performed by Metabolon, Inc. Briefly, the 23 GBM cell lines were cultured in DMEM to 75-90% confluence and harvested one hour after addition of fresh media (Fig. 1C). Two RT-resistant cell lines (U87 and A172) and two RT-sensitive cell lines (KS-1 and U118 MG, Fig. 1D & E) were maintained in cell culture to 75-90% confluence. After 1 h in fresh culture medium, cells were irradiated (8 Gy) and harvested after an additional 2 h incubation. The global metabolic profiles of all the harvested cell samples (4-5 biologic replicates per condition) were determined by Metabolon. Data from this profiling effort is attached a supplementary spreadsheet.

### Metabolic Pathway Analysis

The normalized metabolite intensity levels were z-transformed (i.e. zero mean and unit variance across all cell lines) to enable comparison on the same scale^54^. Metabolites with z-score below −1 or above +1 were assumed to be downregulated or upregulated. Metabolites were then grouped into corresponding pathways based on annotation from Metabolon. A pathway-level up or downregulation score was determined by calculating the ratio of the total number of metabolites that were significantly up or down regulated to the total metabolites measured in that pathway. This was then correlated with RT-resistance score using Pearson’s linear correlation function in MATLAB. Downregulated pathways with significant correlation with RT-resistance (*p* < 0.05) in pre-treatment condition and post-treatment are shown in Fig. 1C and 1E respectively. The pathways with significant correlation (*p* < 0.05) were also found to be significant after Benjamin-Hochberg false discovery rate correction (FDR < 0.1). Significantly correlated pathways were then visualized on a human metabolic network map (Fig. S1H) using the iPath pathway explorer^55^.

### TCGA clinical and molecular data

The Cancer Genome Atlas (TCGA) PanCancer Atlas LGG and GBM cohorts were used for survival analysis and gene expression profiling^56^. For the purposes of the current study, curation of these cases was performed to include only *IDH* wild type primary/untreated samples, WHO grades II-IV. We further excluded cases based on those that were masked (“Do_not_use”) according to the Pan-Cancer Atlas sample quality annotations (http://api.gdc.cancer.gov/data/1a7d7be8-675d-4e60-a105-19d4121bdebf). From the initial 1,118 cases identified in the LGG and GBM project, 235 IDH-wildtype primary tumors were used for further analyses.

Gene expression data (RNA-seq) from the LGG and GBM cohorts was downloaded from cBioPortal (http://www.cbioportal.org/, accessed 9/29/2019). RSEM-normalized expression values were then stratified into low and high expressing groups using a median cutoff. Overall survival was used as the clinical endpoint and survival analytics were obtained from the TCGA Pan-Cancer Clinical Data Resource (TCGA-CDR). Survival curves were generated using the Kaplan-Meier method and significance assessed using the log rank test. Statistical significance was set at *p* < 0.05.

### Statistical methods

Clonogenic survival, γ-H2AX foci formation, comet assay, Ki-67 IHC staining, and metabolite level analysis after MPA and Teriflunomide treatment were analyzed by two-tailed t tests using GraphPad Prism Version 8 with the Holm-Sidak method employed to account for multiple comparisons when appropriate. Tumor volume of GBM xenografts were normalized to 100% at the first day for each group and growth rates were analyzed using a linear mixed effects model. Time to tumor tripling in each group was determined by identifying the earliest day on which it was at least three times as large as on the first day of treatment and then estimated by the Kaplan-Meier method and compared using the log-rank test. Significance threshold was set at *p* < 0.05.

## Supporting information

Mass Spectrometry dataset 1

Mass Spectrometry dataset 2

## Acknowledgments

We thank Rob Mohney, PhD, Ed Karoly PhD and colleagues at Metabolon Inc for their metabolomics profiling of GBM cell lines. We thank Steven Krongenberg for his assistance with illustrations. D.R.W was supported by grants from the American Cancer Society, the Forbes Institute for Cancer Discovery, the NCI (K08CA234416) and the Jones Family Foundation Fund within the Chad Carr Pediatric Brain Tumor Center. W.Z was supported by the Postdoctoral Translational Scholar Program (UL1TR002240) from the Michigan Institute for Clinical & Health Research of University of Michigan. A.J.S was supported by the Rogel Cancer Center Post-doctoral Research Fellowship (G023496). C.J.H was supported by P30DK034933 and F32CA228328. C.A.L. was supported by a 2017 AACR NextGen Grant for Transformative Cancer Research (17-20-01-LYSS) and an ACS Research Scholar Grant (RSG-18-186-01). Metabolomics studies performed at the University of Michigan were supported by NIH grant DK097153. D.R.W. and C.A.L. were received support from UMCCC Core Grant (P30CA046592). T.S.L was supported by NIH grant U01CA216440 and UMCCC Core Grant P30CA46592.

## Authors’ Contributions

D.R.W, and W.Z conceptualized and designed the study. W.Z, YY, J.J.D, A.J.S, K.W.R and D.R.W developed the methods and performed the experiments. L.Z, B.S.N, C.J.H, C.A.L, and D.R.W completed the data acquisition of LC/MS. P.S, S.G.Z, F.G, S. C, D.P, W.Z, Y.Y, and D.R.W analyzed and interpreted the data. W.Z, Y.Y, C.A.L, T. S.L, A.J.S, M.G.C, P.L, and D.R.W wrote, reviewed and revised the paper. M.D, M.A.M, C.K.W, H.S, A.R, J.X, M.G.C, and P.L provided administrative, technical and material supports. C.A.L and D.R.W supervised the study.

