## Supplemental Figures and Figure legend for "Purine metabolism regulates DNA repair and therapy resistance in glioblastoma"

Fig. S1

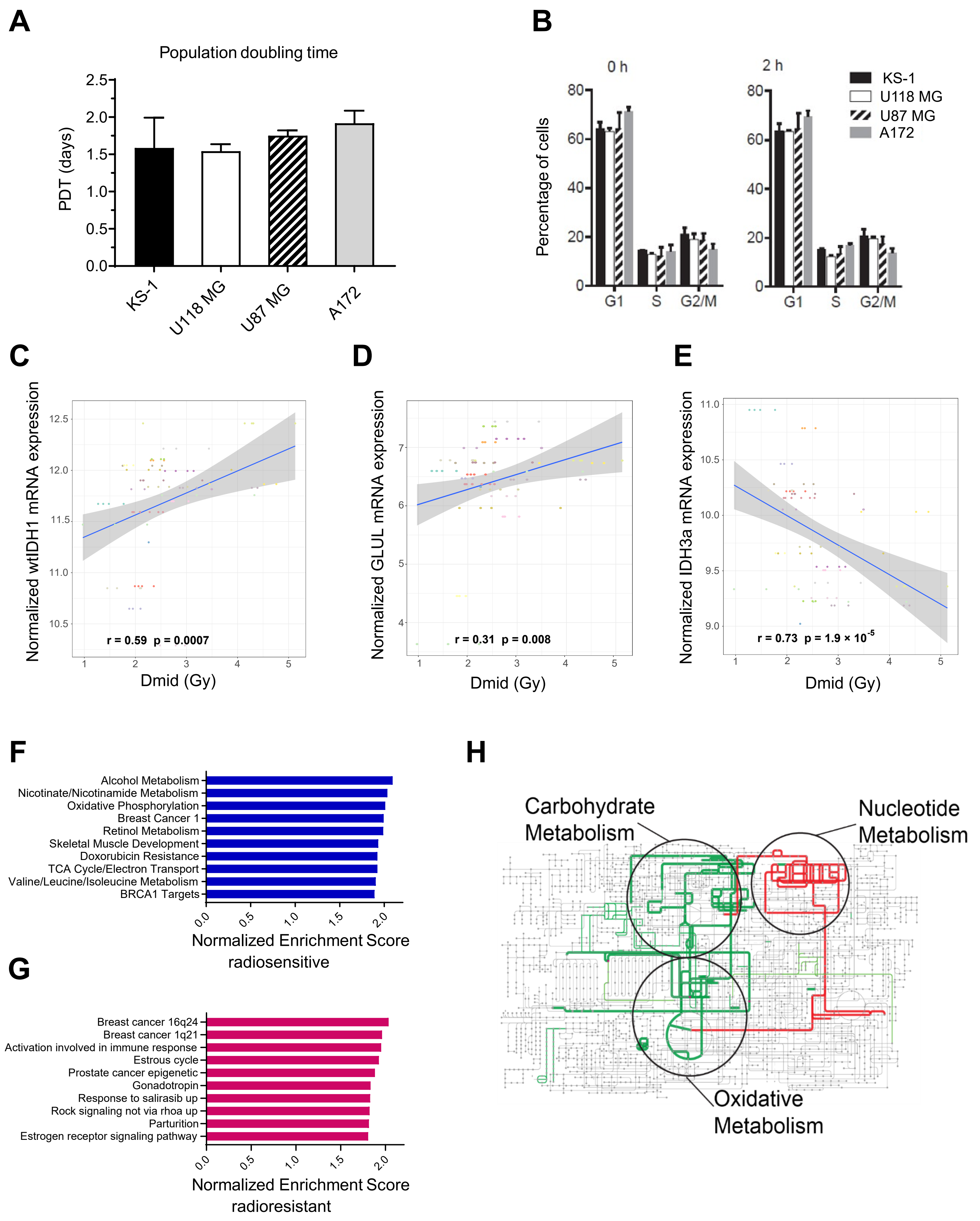
A
Population doubling time
PDT (days)
2.5
2.0
1.5
1.0
0.5
0.0
KS-1
U118 MG
U87 MG
A172
B
0 h
2 h
Percentage of cells
80
60
40
20
0
G1
S
G2/M
KS-1
U118 MG
U87 MG
A172
C
Normalized wtIDH1 mRNA expression
r = 0.59 p = 0.0007
Dmid (Gy)
D
Normalized GLUL mRNA expression
r = 0.31 p = 0.008
Dmid (Gy)
E
Normalized IDH3a mRNA expression
r = 0.73 p = 1.9 × 10^-5
Dmid (Gy)
F
Alcohol Metabolism
Nicotinate/Nicotinamide Metabolism
Oxidative Phosphorylation
Breast Cancer 1
Retinol Metabolism
Skeletal Muscle Development
Doxorubicin Resistance
TCA Cycle/Electron Transport
Valine/Leucine/Isoleucine Metabolism
BRCA1 Targets
Normalized Enrichment Score
radiosensitive
G
Breast cancer 16q24
Breast cancer 1q21
Activation involved in immune response
Estrous cycle
Prostate cancer epigenetic
Gonadotropin
Response to salirasib up
Rock signaling not via rhoa up
Parturition
Estrogen receptor signaling pathway
Normalized Enrichment Score
radioresistant
H
Carbohydrate Metabolism
Nucleotide Metabolism
Oxidative Metabolism

**Figure S1. Correlates with GBM RT-sensitivity**

**(A)** Indicated GBM cell lines were plated, and cell number were counted 3 to 8 days later. **(B)** RT-resistant (U87 MG and A172) and RT-sensitive (KS-1 and U118 MG) GBM cell lines were irradiated with 8 Gy, and cells were harvested for flow cytometry cell cycle analysis 2 h post-RT. **(C-E)** Correlation analysis between *IDH1*, *IDH3*a, *GLUL* transcript levels and RT-sensitivity of GBMs. **(F&G)** GSEA analysis of pathway correlates of RT-sensitivity and RT-resistance. **(H)** Pathways with downregulated metabolites that are correlated with RT-sensitivity are shown on a network map of human metabolism (Pearson's correlation $p < 0.08$). Pathways with negative and positive correlation with RT-sensitivity are shown in green and red respectively.

Fig. S2

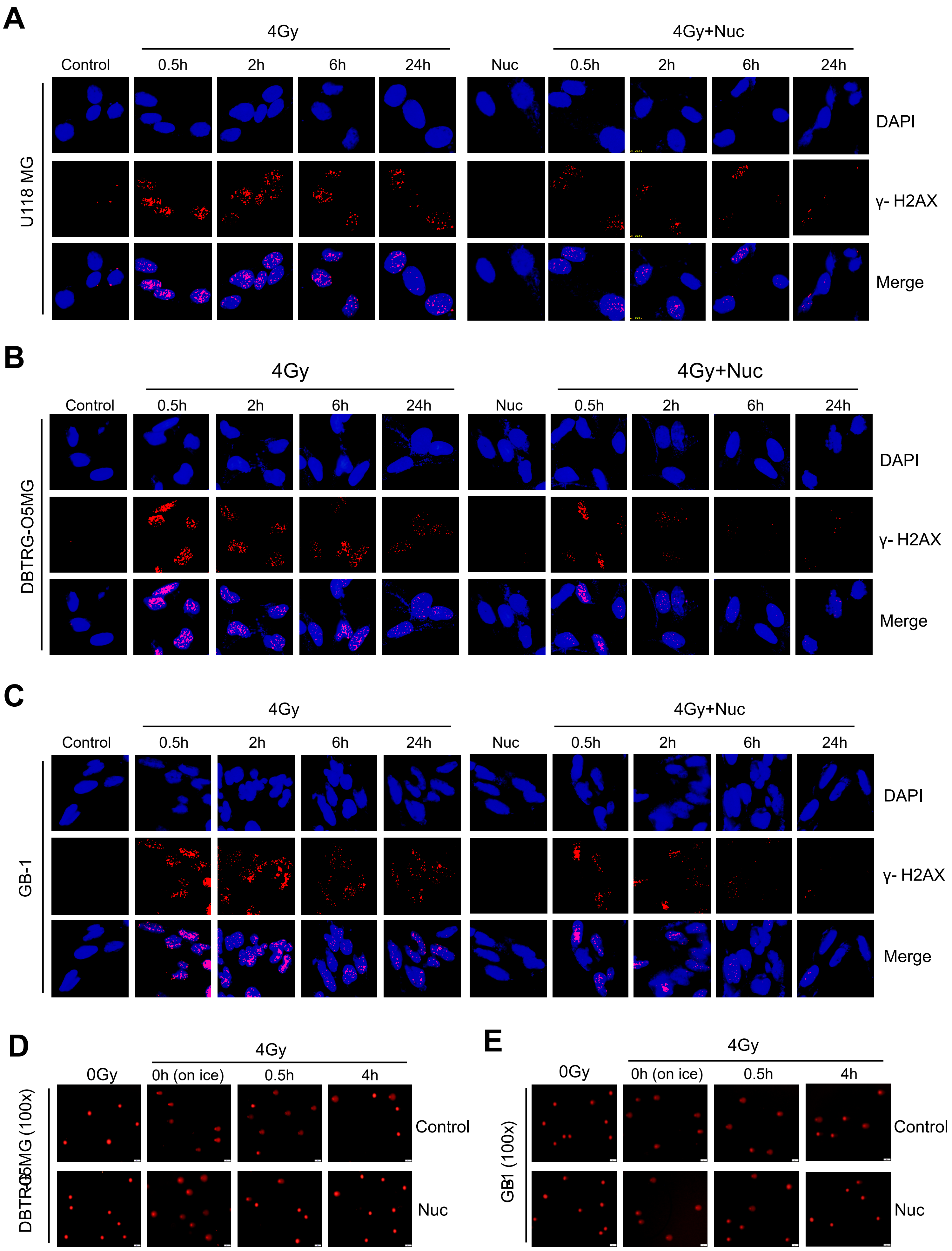
A
4Gy
4Gy+Nuc
Control
0.5h
2h
6h
24h
Nuc
0.5h
2h
6h
24h
U118 MG
DAPI
γ- H2AX
Merge
B
4Gy
4Gy+Nuc
Control
0.5h
2h
6h
24h
Nuc
0.5h
2h
6h
24h
DBTRG-O5MG
DAPI
γ- H2AX
Merge
C
4Gy
4Gy+Nuc
Control
0.5h
2h
6h
24h
Nuc
0.5h
2h
6h
24h
GB-1
DAPI
γ- H2AX
Merge
D
4Gy
0Gy
0h (on ice)
0.5h
4h
DBTRG-O5MG (100x)
Control
Nuc
E
4Gy
0Gy
0h (on ice)
0.5h
4h
GB1 (100x)
Control
Nuc

**Figure S2. Exogenous nucleosides protect RT-sensitive GBMs from RT**

**(A-C)** Representative images of γ-H2AX foci staining in U118 MG (corresponding to Fig. 2E), DBTRG-05MG (corresponding to Fig. 2F) and GB-1 (corresponding to Fig. 2G) cells. Cells were treated with nucleoside pools, irradiated and fixed for γ-H2AX foci staining at indicated time point post-RT. **(D&E)** Representative images of alkaline comet assay in DBTRG-05MG (corresponding to Fig. 2H) and GB-1 (corresponding to Fig. 2I) cells. DBTRG-05MG or GB-1 cells were plated, treated with exogenous pooled nucleosides, and analyzed by alkaline comet assay at the indicated time points.

Fig. S3

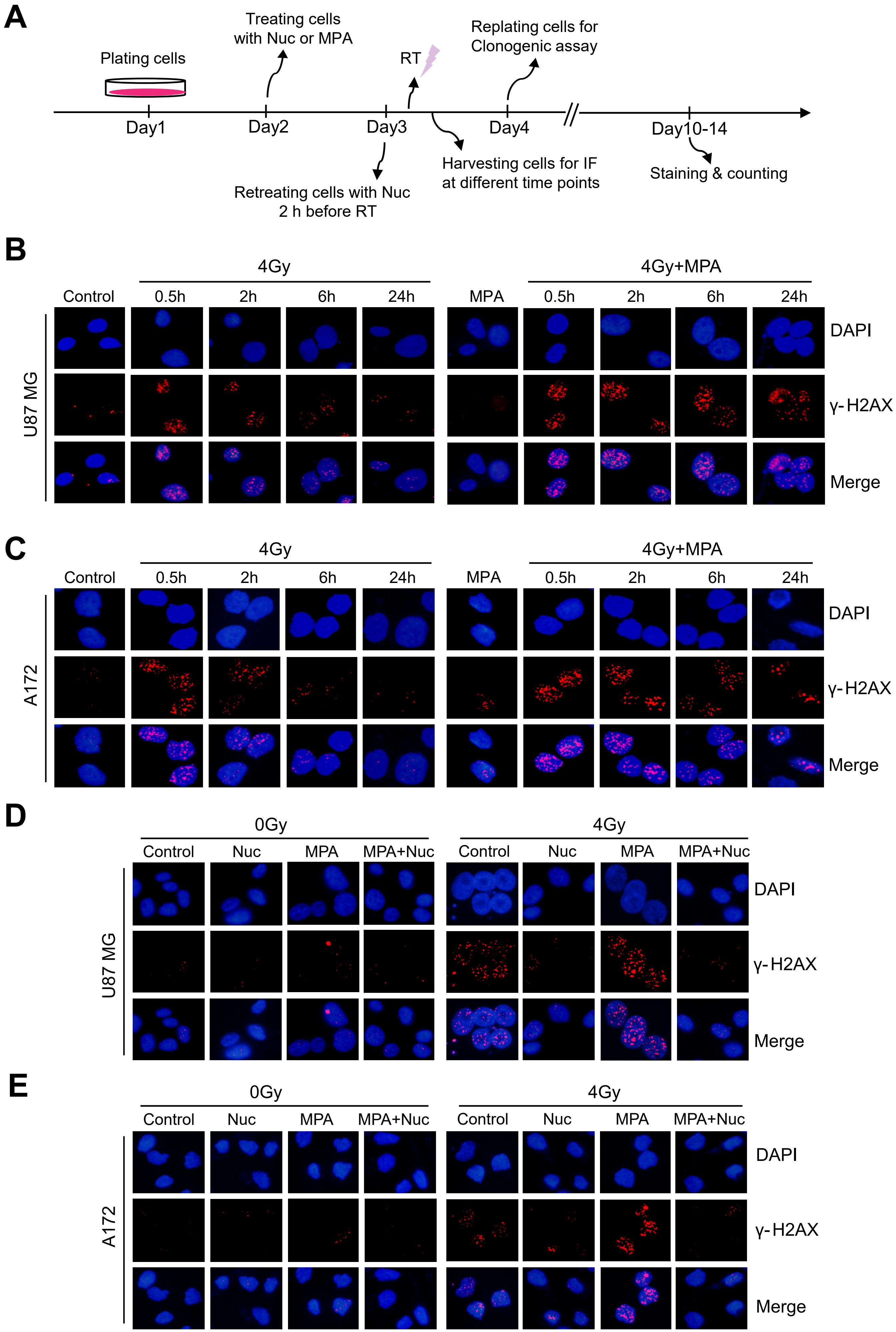

A
Plating cells
Treating cells with Nuc or MPA
RT
Replating cells for Clonogenic assay
Day1
Day2
Day3
Day4
Day10-14
Retreating cells with Nuc 2 h before RT
Harvesting cells for IF at different time points
Staining & counting
B
4Gy
4Gy+MPA
Control
0.5h
2h
6h
24h
MPA
0.5h
2h
6h
24h
U87 MG
DAPI
γ-H2AX
Merge
C
4Gy
4Gy+MPA
Control
0.5h
2h
6h
24h
MPA
0.5h
2h
6h
24h
A172
DAPI
γ-H2AX
Merge
D
0Gy
4Gy
Control
Nuc
MPA
MPA+Nuc
Control
Nuc
MPA
MPA+Nuc
U87 MG
DAPI
γ-H2AX
Merge
E
0Gy
4Gy
Control
Nuc
MPA
MPA+Nuc
Control
Nuc
MPA
MPA+Nuc
A172
DAPI
γ-H2AX
Merge

**Figure S3. Inhibiting *de novo* purine synthesis impairs DNA repair and radiosensitizes RT-resistant GBM cells**

**(A)** A schematic timeline of treatment administrated in RT-resistant cell lines. Cells were treated with pooled nucleosides and/or MPA for 24 h, and retreated with pooled nucleosides 2 h before RT (4 Gy), followed by clonogenic assay or $\gamma$-H2AX foci staining. **(B&C)** Representative images of $\gamma$-H2AX foci staining in U87 MG (corresponding to Fig. 3G), and A172 (corresponding to Fig. 3H) cells. Cells were treated with different doses of MPA for 24 h and irradiated with 4 Gy, followed by $\gamma$-H2AX foci IF staining. (**D&E**) Representative images of $\gamma$-H2AX foci IF staining in U87 MG (corresponding to Fig. 3I), and A172 (corresponding to Fig. 3J) cells. Cells were treated as described above, followed by $\gamma$-H2AX foci staining 6 h post-RT.

### Fig. S4

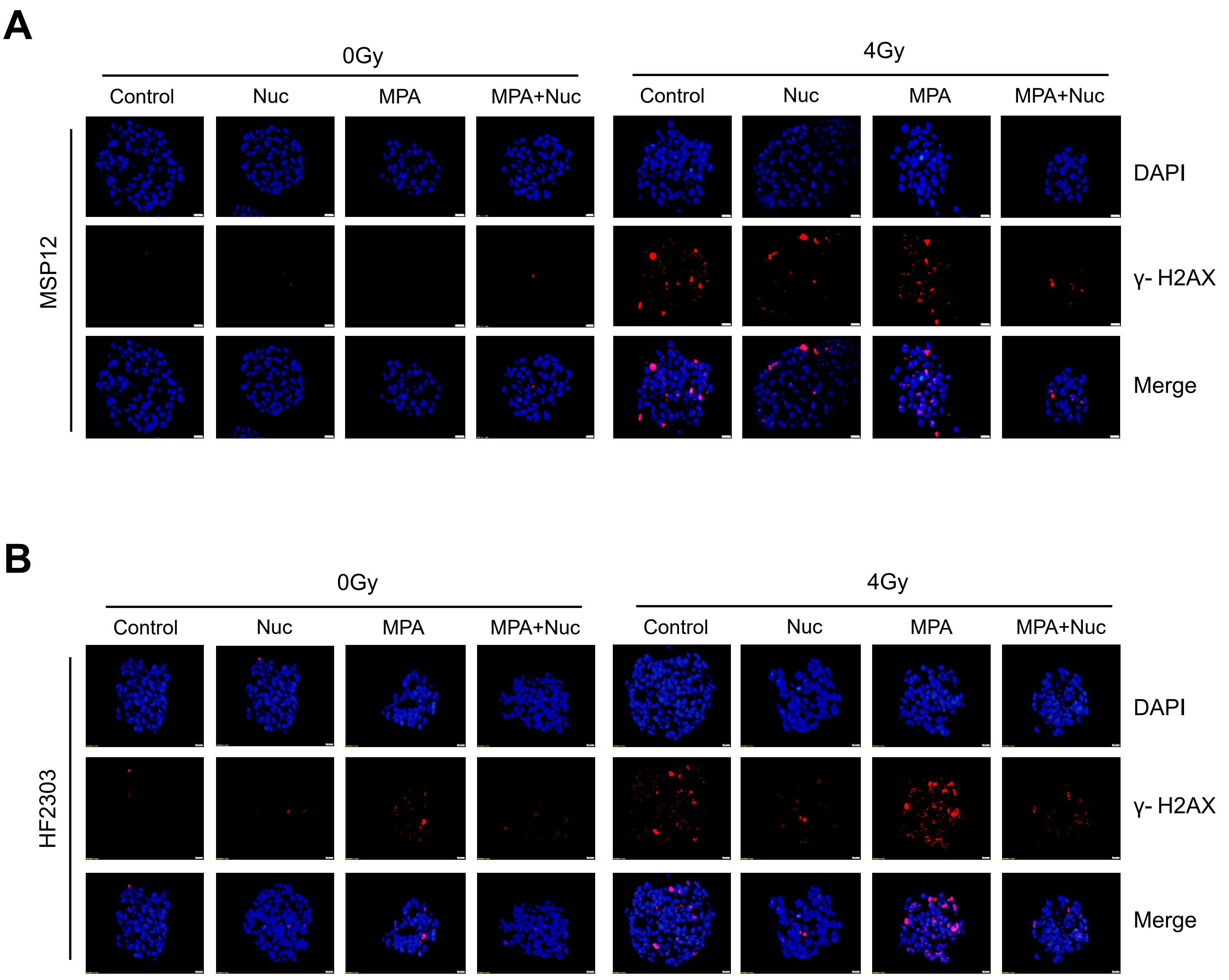
A
0Gy
4Gy
Control
Nuc
MPA
MPA+Nuc
Control
Nuc
MPA
MPA+Nuc
MSP12
DAPI
γ- H2AX
Merge
B
0Gy
4Gy
Control
Nuc
MPA
MPA+Nuc
Control
Nuc
MPA
MPA+Nuc
HF2303
DAPI
γ- H2AX
Merge

**Figure S4. Inhibiting *de novo* purine synthesis radiosensitizes primary patient-derived GBM neurospheres in a nucleoside-dependent fashion**

**(A&B**) Representative images of γ-H2AX foci IF staining in MSP12 (corresponding to Fig. 4C), and HF2303 (corresponding to Fig. 4D) cells. Cells were treated by indicated compounds and then irradiated, followed by γ-H2AX foci staining 6 h post-RT.

Fig. S5

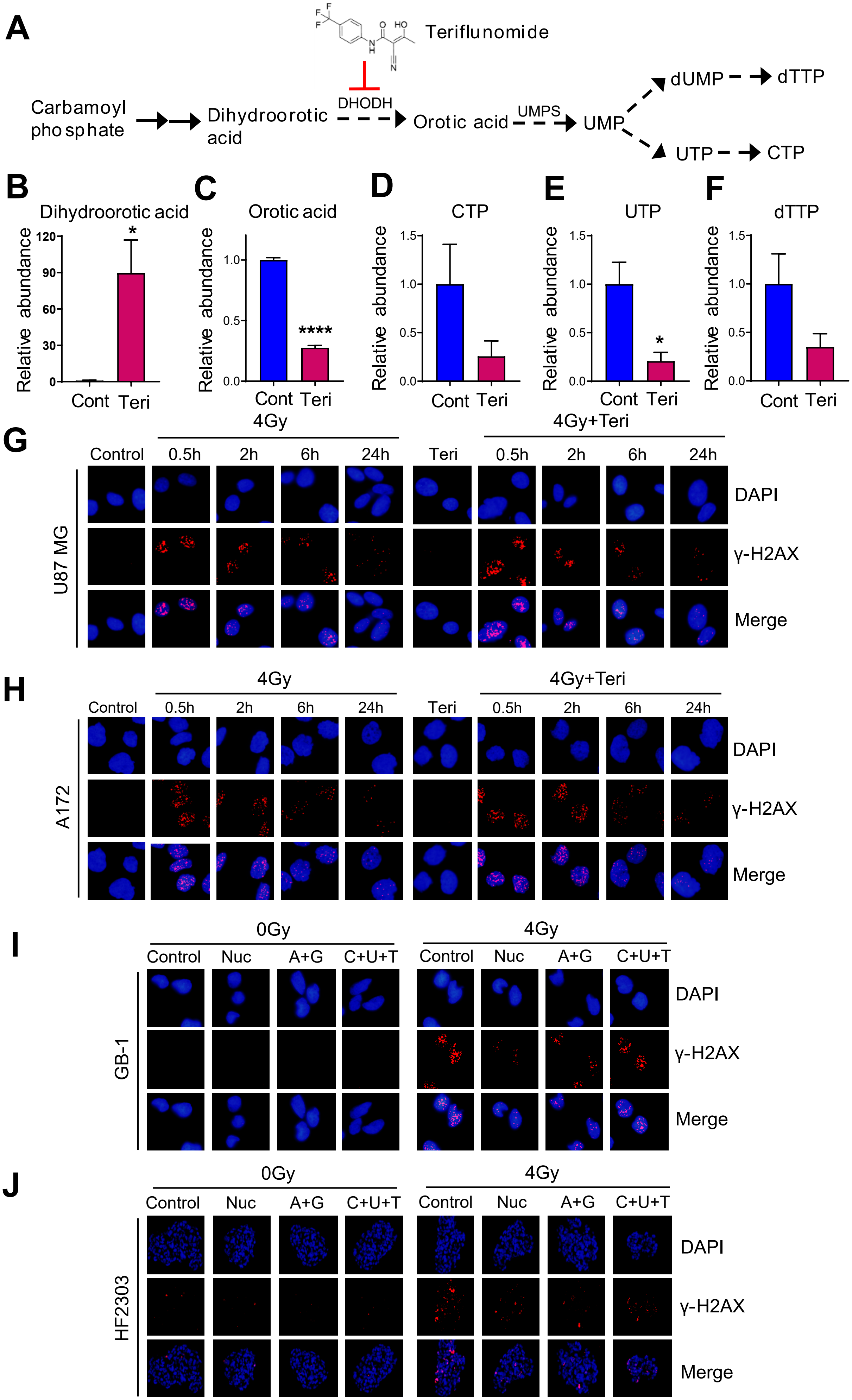
A
Teriflunomide
Carbamoyl phosphate
Dihydroorotic acid
DHODH
Orotic acid
UMPS
UMP
dUMP
dTTP
UTP
CTP
B
Dihydroorotic acid
Relative abundance
120
90
60
30
0
*
Cont
Teri
C
Orotic acid
Relative abundance
1.0
0.5
0.0
****
Cont
Teri
D
CTP
Relative abundance
1.5
1.0
0.5
0.0
Cont
Teri
E
UTP
Relative abundance
1.5
1.0
0.5
0.0
*
Cont
Teri
F
dTTP
Relative abundance
1.5
1.0
0.5
0.0
Cont
Teri
G
4Gy
4Gy+Teri
Control
0.5h
2h
6h
24h
Teri
0.5h
2h
6h
24h
U87 MG
DAPI
γ-H2AX
Merge
H
4Gy
4Gy+Teri
Control
0.5h
2h
6h
24h
Teri
0.5h
2h
6h
24h
A172
DAPI
γ-H2AX
Merge
I
0Gy
4Gy
Control
Nuc
A+G
C+U+T
Control
Nuc
A+G
C+U+T
GB-1
DAPI
γ-H2AX
Merge
J
0Gy
4Gy
Control
Nuc
A+G
C+U+T
Control
Nuc
A+G
C+U+T
HF2303
DAPI
γ-H2AX
Merge

**Figure S5. Modulating pyrimidine pools has minimal effects on DNA repair and RT-resistance in GBM**

**(A)** A schematic diagram of *de no* pyrimidine synthesis. DHODH: dihydroorotate dehydrogenase; UMPS: uridine monophosphate synthetase; UMP: uridine monophosphate; dUMP: deoxyuridine monophosphate; dTTP: deoxythymidine triphosphate; UTP: uridine triphosphate; CTP: cytidine triphosphate. **(B-F)** U87 MG cells were treated with 20 μM teriflunomide for 24 h, and then harvested for targeted LC/MS assay. *, $p < 0.05$; ****, $p < 0.0001$. **(G&H)** Representative images of γ-H2AX foci IF staining in U87 MG (corresponding to Fig. 5C), and A172 (corresponding to Fig. 5D) cells. Cells were treated by teriflunomide (20 μM) for 24 h, and irradiated (4 Gy), and harvested at different time points post-RT for γ-H2AX foci staining. **(I&J)** Representative images of γ-H2AX foci in GB-1 (corresponding to Fig. 5E), and HF2303 (corresponding to Fig. 5F) cells. Cells were treated by indicated compounds and then irradiated, followed by γ-H2AX foci staining 6 h post-RT.

**Fig. S6**

**A**

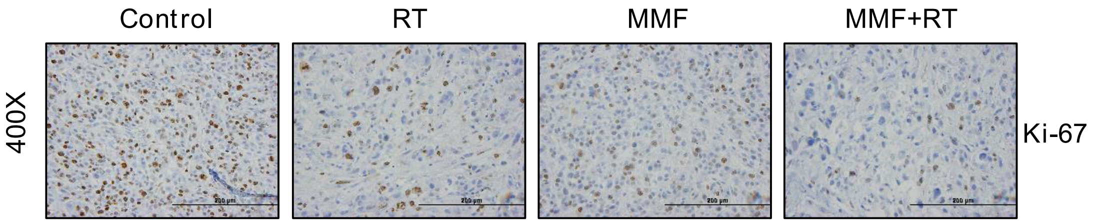
Control
RT
MMF
MMF+RT
400X
Ki-67

**B**

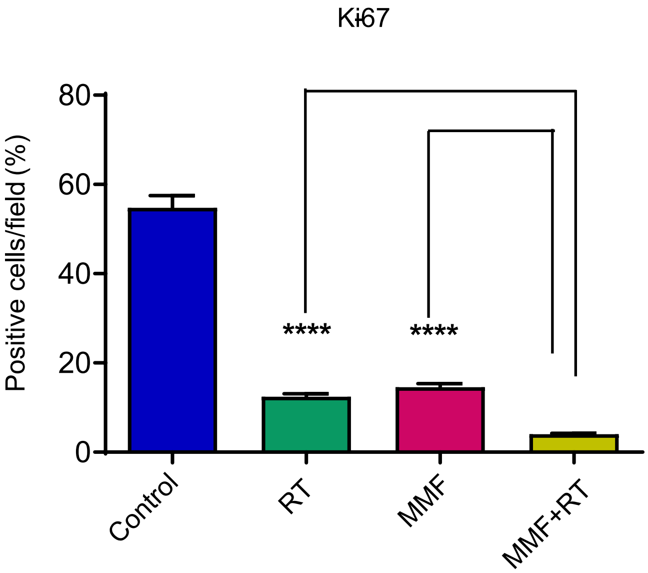
Ki67
Positive cells/field (%)
80
60
40
20
0
****
****
Control
RT
MMF
MMF+RT

**Figure S6. MMF augments RT efficacy against immortalized GBM xenografts**

**(A&B)** Representative images and analysis of Ki-67 expression in four arms of treatment. Staining quantification: positively stained cells were counted out of 5 different fields for each tumor, with an average from 3 independent tumors derived from 3 mice per group. ****, $p < 0.0001$.

### Fig. S7

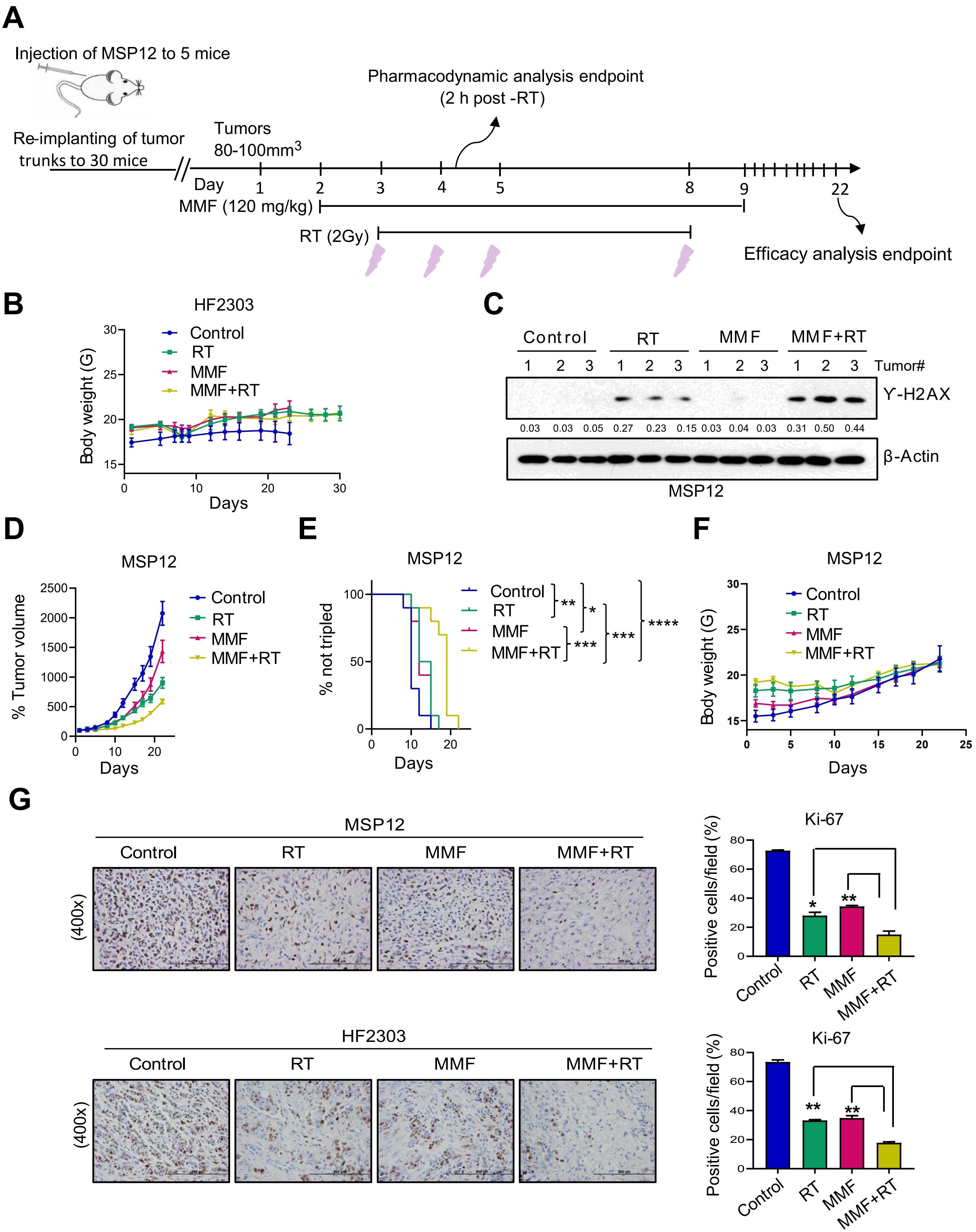
A
Injection of MSP12 to 5 mice
Re-implanting of tumor trunks to 30 mice
Tumors 80-100mm3
Pharmacodynamic analysis endpoint (2 h post -RT)
Day 1 2 3 4 5 8 9 22
MMF (120 mg/kg)
RT (2Gy)
Efficacy analysis endpoint
B
HF2303
Control
RT
MMF
MMF+RT
Body weight (G)
Days
C
Control RT MMF MMF+RT
1 2 3 1 2 3 1 2 3 1 2 3 Tumor#
γ-H2AX
0.03 0.03 0.05 0.27 0.23 0.15 0.03 0.04 0.03 0.31 0.50 0.44
β-Actin
MSP12
D
MSP12
% Tumor volume
Days
E
MSP12
% not tripled
Days
**
*
***
***
****
F
MSP12
Body weight (G)
Days
G
MSP12
HF2303
Control RT MMF MMF+RT
(400x)
Ki-67
Positive cells/field (%)
Control
RT
MMF
MMF+RT

**Figure S7. MMF augments RT efficacy against patient-derived GBM xenografts**

**(A)** MSP12 xenograft GBM model was established and treated as described in Material and Methods. **(B)** Body weight curve of HF2303 GBM model. Body weight was measured during the treatment and plotted (mean ± SEM). **(C)** Flash-frozen tumors harvested 2 h post-RT in biological group were analyzed by immunoblotting. The bands were quantified using Image J software and the quantified numbers were labeled under each band. **(D)** Tumor volumes for the indicated treatment subgroups of the efficacy group are normalized to the individual tumor sizes defined on day 1. Error bars indicate the SEM from 10 tumors from 5 mice per group. **(E)** Kaplan-Meier estimates of time to tumor tripling. *, $p < 0.05$; **, $p < 0.01$; ***, $p < 0.001$; ****, $p < 0.0001$. (**F**) Body weight curve of MSP12 GBM model. Body weight was measured during the treatment and plotted (mean ± SEM). **(G)** Representative images and analysis of Ki-67 expression in HF2303 and MSP12 xenograft models. *, $p < 0.05$; **, $p < 0.01$; mean ± SEM.

**Table S1 Details of the GBM cell lines sources**

| Cell line | Company/Source | Catalog No. |
|---|---|---|
| U87 MG | ATCC | HTB-14 |
| A172 | ATCC | CRL-1620 |
| DKMG | DSMZ | ACC 277 |
| 8MGBA | DSMZ | ACC 432 |
| 42MGBA | DSMZ | ACC 431 |
| KNS-60 | JCRB | IFO50357 |
| AM-38 | JCRB | IFO50492 |
| M059K | ATCC | CRL-2365 |
| KNS-81 | JCRB | IFO50359 |
| YKG-1 | JCRB | JCRB0746 |
| U138 | ATCC | HTB-16 |
| T98G | ATCC | CRL-1690 |
| LN18 | ATCC | CRL-2610 |
| SNB19 | DSMZ | ACC 325 |
| GOS3 | DSMZ | ACC 408 |
| LN229 | ATCC | CRL-2611 |
| GMS10 | DSMZ | ACC 405 |
| GB-1 | JCRB | IFO50489 |
| U118 MG | ATCC | HTB-15 |
| SW1783 | ATCC | HTB-13 |

| | | |
|---|---|---|
| KS-1 | JCRB | IFO50436 |
| DBTRG-05MG | ATCC | CRL-2020 |

ATCC: American Type Culture Collection

DSMZ: Leibniz Institute DSMZ-German Collection of Microorganisms and Cell Cultures

JCRB: Japanese Collection of Research Bioresources Cell Bank
